# Cytokinin activity of *N*^6^-benzyladenine derivatives assayed by interaction with the receptors *in planta, in vitro*, and *in silico*

**DOI:** 10.1101/241281

**Authors:** Ekaterina M. Savelieva, Vladimir E. Oslovsky, Dmitry S. Karlov, Nikolay N. Kurochkin, Irina A. Getman, Sergey N. Lomin, Georgy V. Sidorov, Sergey N. Mikhailov, Dmitry I. Osolodkin, Georgy A. Romanov

## Abstract

Biological effects of hormones in both plants and animals are based on high-affrnity interaction with cognate receptors resulting in their activation. The signal of cytokinins, classical plant hormones, is perceived in *Arabidopsis* by three homologous membrane receptors: AHK2, AHK3, and CRE1/AHK4. To study the cytokinin–receptor interaction, we used 25 derivatives of potent cytokinin *N*^6^-benzyladenine (BA) with substituents in the purine heterocycle and/or in the side chain. The study was focused primarily on individual cytokinin receptors from *Arabidopsis*. The main *in planta* assay system was based on *Arabidopsis* double mutants retaining only one isoform of cytokinin receptors and harboring cytokinin-sensitive reporter gene. Classical cytokinin biotest with *Amaranthus* seedlings was used as an additional biotest. In parallel, the binding of ligands to individual cytokinin receptors was assessed in the *in vitro* test system. Quantitative comparison of results of different assays confirmed the partial similarity of ligand-binding properties of receptor isoforms. Substituents at positions 8 and 9 of adenine moiety, elongated linker up to 4 methylene units, replacement of *N*^6^ by sulfur or oxygen, resulted in suppression of cytokinin activity of the derivative towards all receptors. Introduction of a halogen into position 2 of adenine moiety, on the contrary, often increased the ligand activity, especially toward AHK3. Features both common and distinctive of cytokinin receptors in *Arabidopsis* and *Amaranthus* were revealed, highlighting species specificity of the cytokinin perception apparatus. Correlations between extent of compound binding to a receptor *in vitro* and its ability to activate the same receptor *in planta* were evaluated for each AHK protein. Interaction patterns between individual receptors and ligands were rationalized by structure analysis and molecular docking in sensory modules of AHK receptors. The best correlation between docking scores and specific binding was observed for AHK3. In addition, receptor-specific ligands have been discovered with unique properties to predominantly activate or block distinct cytokinin receptors. These ligands are promising for practical application and as molecular tools in the study of the cytokinin perception by plant cells.

**Graphical abstract:** 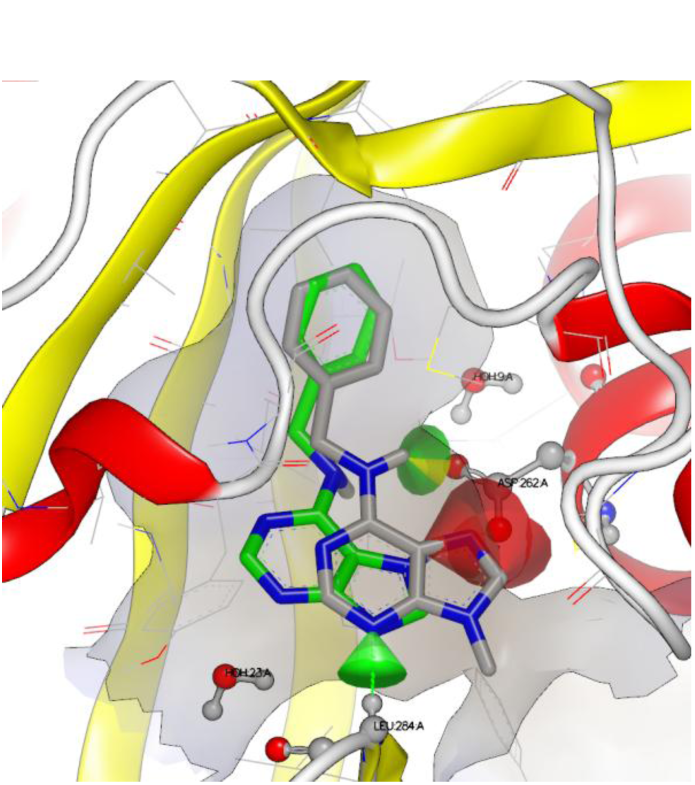

Individual cytokinin receptors from *Arabidopsis* were assayed *in planta*, *in vitro* and *in silico* with 25 different 6-benzyladenine derivatives, new receptor-specific cytokinins were revealed.

## 1. Introduction

Cytokinins are classical phytohormones that play an important role at all stages of plant growth and development. They stimulate cell division, regulate shoot and root growth, reduce apical dominance, promote chloroplast differentiation, delay leaf senescence, etc. (Sakakibara, 2006; Romanov, 2009). Common cytokinins are derivatives of adenine or phenylurea. Phenylurea derivatives are synthetic cytokinins such as thidiazuron, whereas adenine derivatives can be both natural and synthetic hormones. Natural cytokinins may, in turn, be divided into isoprenoid ones, which include isopentenyladenine, zeatins and dihydrozeatin, and aromatic ones, for example, *N*^6^-benzyladenine and topolins.

Cytokinin signal is perceived through transmembrane receptors having a complex multidomain structure and histidine kinase activity (Heyl et al., 2012; Lomin et al., 2012; Steklov et al., 2013). Further signal transmission occurs by multistep phosphorelay, which results in the phosphorylation of transcription factors (B-type response regulators) in the cell nucleus (Kieber and Schaller, 2014) followed by changes in the activity (mainly upregulation) of primary response genes (Rashotte et al., 2003; Brenner et al., 2005). In plants, cytokinin receptors are usually represented by small families of a few isoforms. In particular, *Arabidopsis* has three receptor isoforms: AHK2, AHK3 and CRE1/AHK4 (Kakimoto, 2003). They are structurally similar and functionally complement each other to a high extent. However, due to some differences in structure, they differ in ligand specificity and are unequal in a number of physiological processes (Stolz et al., 2011; Lomin et al., 2012).

Cytokinin receptors are large transmembrane proteins, that greatly hampers their crystallization. Therefore, a complete 3D structure has not yet been solved for any of cytokinin receptors. So far, only structure of CRE1/AHK4 sensory domain complexed with various cytokinins was reported (Hothorn et al., 2011). It has been shown by biochemical (Romanov et al., 2006) and structural (Hothorn et al., 2011) methods that CRE1/AHK4 receptor recognizes both natural and synthetic cytokinins with the same binding site. Thus, extension of spectrum of ligands interacting with cytokinin receptors may be done by systematic analogue synthesis. Use of a wide range of ligands allows to reveal the features of their structure that are important for interaction with the sensory domain, and to characterize its ligand-binding properties more accurately.

For our study, we have selected a range of derivatives of a potent cytokinin *N*^6^-benzyladenine (BA), an aromatic cytokinin found in a number of plant species (Strnad, 1997). For decades, BA has been widely used in tissue culture as an inducer of cell division and shoot differentiation. High activity of BA *in planta* is attributed to its resistance to enzymatic degradation and modification (Laloue and Fox, 1989). However, BA is not a perfect plant growth stimulator since it exerts not only positive, but also negative effects upon plant treatment, in particular, inhibits root growth.

Biological effects of BA derivatives have been extensively studied. Synthesis and testing of these compounds in plant assay systems began soon upon the first cytokinin discovery (Miller et al., 1955), but results of early studies conducted before receptor identification were rather ambiguous (Matsubara, 1990). Upon the first cytokinin receptor discovery (Inoue et al., 2001; Suzuki et al., 2001), specific assay systems based on ligand-receptor interaction *in vitro* and *in vivo* (Spíchal et al., 2004; Romanov et al., 2005; Romanov and Lomin, 2009; Stolz et al., 2011; Lomin et al., 2015) were developed. Among BA derivatives studied in these assays, many represented BA or BA ribosides with substituents in phenyl ring (Doležal et al., 2006, 2007; Szücová et al., 2009; Podlešáková et al., 2012; Plíhal et al., 2013) alone or together with substituents at position 9 of adenine moiety (Szücová et al., 2009; Podlešáková et al., 2012; Plíhal et al., 2013). In addition, 9-substituted 2-chloro-BA ribosides (Vylíčilová et al., 2016) and 8-substituted BA (Zahajská et al., 2017) were recently studied in various assay systems. Numerous synthetic derivatives of BA were shown to possess genuine cytokinin activity. Several BA-related compounds were found to have anticytokinin (Spíchal et al., 2009; Nisler et al., 2010; Krivosheev et al., 2012) or anticancer (Kolyachkina et al., 2011; Voller et al., 2017 and refs therein) activities or inhibitory effect on cytokinin oxidase/dehydrogenase (Zatloukal et al., 2008). Contrary to unmodified BA, its methoxy derivative topolin demonstrated at nanomolar concentration a positive effect on lateral root branching and leaf emerging (Podlešáková et al., 2012).

In this study we used compounds differing from BA by (i) various substitutions at different positions of the purine heterocycle; (ii) length of the linker between purine and phenyl moieties; (iii) substitutions of nitrogen in linker for oxygen or sulfur, and (iv) combinations of these modifications. Special attention was given to differences in ligand specificity between *Arabidopsis* receptors (AHK2, AHK3, and CRE1/AHK4). Upon assessment of cytokinin activity for BA derivatives using *in vivo* assay with *Arabidopsis* receptor double mutants (Stolz et al., 2011) and *in vitro* binding to individual receptors (Romanov et al., 2005; Lomin et al., 2015), we found compounds specific to individual isoforms of the *Arabidopsis* cytokinin receptors. New traits of ligand structure important for specific binding to receptors were revealed. These data supported with molecular modeling shed light on structural features of receptor sensor modules providing their competence in cytokinin binding as well as differences in ligand preference.

## 2. Results

### 2.1. *Cytokinin activity of BA analogues in the biotest with* Arabidopsis *double receptor mutants*

To characterize individual cytokinin receptors, we used three double receptor mutants of *Arabidopsis*, each retaining a single receptor, along with WT plants. Activation of the cytokinin-sensitive reporter *P_ARR_*_5_*:GUS* construct introgressed into genomes of double mutants and WT plants was assessed. BA derivatives (Table 1; Supplementary Table 1) were tested along with BA serving as a positive control. Activity of BA was taken for 100%. Activity of other compounds was determined in % relative to BA activity.

**Table 1.** Studied BA derivatives a.

| Compound | Scaffold | R | R<sub>1</sub> | L | Ar |
| --- | --- | --- | --- | --- | --- |
| **1** (BA) | A | H | H | NHCH<sub>2</sub> | phenyl |
| **2** | A | methyl | H | NHCH<sub>2</sub> | phenyl |
| **3** | A | ethyl | H | NHCH<sub>2</sub> | phenyl |
| **4** | A | isopropyl | H | NHCH<sub>2</sub> | phenyl |
| **5** | B (X = O) | methyl | H | NHCH<sub>2</sub> | phenyl |
| **6** | B (X = O) | H | H | NHCH<sub>2</sub> | phenyl |
| **7** | B (X = S) | H | H | NHCH<sub>2</sub> | phenyl |
| **8** | B (X = S) | H | F | NHCH<sub>2</sub> | phenyl |
| **9** | B (X = S) | H | Cl | NHCH<sub>2</sub> | phenyl |
| **10** | A | H | Cl | NHCH<sub>2</sub> | phenyl |
| **11** | A | H | F | NHCH<sub>2</sub> | phenyl |
| **12** | A | H | Cl | NHCH<sub>2</sub> | 3-fluorophenyl |
| **13** | A | H | NH<sub>2</sub> | NHCH<sub>2</sub> | phenyl |
| **14** | A | H | H | OCH<sub>2</sub> | phenyl |
| **15** | A | H | H | SCH<sub>2</sub> | phenyl |
| **16** | A | H | NH<sub>2</sub> | OCH<sub>2</sub> | phenyl |
| **17** | A | H | NH<sub>2</sub> | O(CH<sub>2</sub>)<sub>2</sub> | phenyl |
| **18** | A | H | NH<sub>2</sub> | O(CH<sub>2</sub>)<sub>3</sub> | phenyl |
| **19** | A | H | H | NH | phenyl |
| **20** | A | H | H | NH(CH<sub>2</sub>)<sub>2</sub> | phenyl |
| **21** | A | H | H | NH(CH<sub>2</sub>)<sub>3</sub> | phenyl |
| **22** | A | H | H | NH(CH<sub>2</sub>)<sub>4</sub> | phenyl |
| **23** | A | H | Cl | NHCH<sub>2</sub> | 2-furyl |
| **24** | A | H | Cl | NHCHMe | phenyl |
| **25** | A | H | H | NHCO | phenyl |
| **26** | A | H | H | NHCH<sub>2</sub> | 2-naphthyl |

<sup>a</sup> In the scheme, R, R<sub>1</sub> – substituents; L – linker; Ar – aromatic residue.

Based on results of the biotest with *Arabidopsis* mutants (Table 2), we subdivided all the tested compounds (at 1 μM concentration) into three conventional groups: low-, medium- and high-active. The first group included compounds whose cytokinin activity did not exceed 30%, the second comprised derivatives with activity from 30 to 80%, and the third group exhibited the activity above 80% of the BA activity. Compounds bearing a substituent at position 9 of the purine moiety (**2**–**5**) show low cytokinin activity in almost all variants of the experiment, irrespective of substituent nature, compound concentration and receptor isoform. Compounds with a monosubstituent at *C*8 (**6** and **7**) also showed low cytokinin activity in mutant clones. However, structurally similar disubstituted compounds **8** and **9** with a halogen atom at position 2 displayed moderate cytokinin activity in all variants of the experiment. The activity raise owing to halogenation of *C*2 was particularly noticeable in lines with functional AHK3 receptor. Other *C*2-halogenated BA derivatives (**10**–**12**, **23**, **24**) also demonstrated their greatest activity in plants harboring this receptor. Racemic compound **24** was receptor-specific, exhibiting cytokinin activity only in plants retaining AHK3 receptor.

**Table 2.** Cytokinin activity of BA derivatives in biotest with double receptor mutants of Parr5:*GUS Arabidopsis*^a^.

| Compound | Relative cytokinin activity (%) in *Arabidopsis* lines with functioning receptors: |  |  |  |
| --- | --- | --- | --- | --- |
|  | AHK2-4 | AHK2 | AHK3 | CRE1/AHK4 |
| 1 | 100 | 100 | 100 | 100 |
| 2 | 6.7 ± 1.0 | 18 ± 1 | 7.0 ± 1.1 | 2.7 ± 0.9 |
| 3 | 17 ± 1 | 5.7 ± 2.4 | 6.9 ± 2.9 | 7.3 ± 0.4 |
| 4 | 15 ± 1 | 10 ± 1 | 7.9 ± 0.9 | 5.6 ± 0.5 |
| 5 | 7.2 ± 0.3 | 11.1 ± 0.2 | 8.1 ± 0.4 | 8.4 ± 0.1 |
| 6 | 40 ± 7 | 14 ± 2 | 4.3 ± 0.1 | 5.4 ± 0.5 |
| 7 | 38 ± 1 | 27 ± 1 | 2.4 ± 0.2 | 17 ± 1 |
| 8 | 39 ± 1 | 27 ± 2 | 44 ± 1 | 37 ± 1 |
| 9 | 46 ± 1 | 29 ± 2 | 31 ± 1 | 34 ± 1 |
| 10 | 91 ± 5 | 73 ± 3 | 95 ± 1 | 64 ± 5 |
| 11 | 131 ± 1 | 101 ± 4 | 134 ± 12 | 59 ± 11 |
| 12 | 84 ± 6 | 83 ± 5 | 144 ± 3 | 83 ± 1 |
| 13 | 68 ± 1 | 41 ± 2 | 52 ± 2 | 93 ± 9 |
| 14 | 31 ± 1 | 68 ± 1 | 64 ± 2 | 48 ± 1 |
| 15 | 12.4 ± 0.2 | 14.6 ± 0.2 | 20 ± 1 | 4.8 ± 0.6 |
| 16 | 7.5 ± 0.1 | 10.7 ± 0.3 | 6.1 ± 0.7 | 2.8 ± 0.5 |
| 17 | 5.0 ± 0.5 | 5.8 ± 0.1 | 10 ± 1 | 1.0 ± 0.6 |
| 18 | 4.0 ± 0.6 | 3.5 ± 2.1 | 5.6 ± 0.4 | 0.7 ± 0.4 |
| 19 | 126 ± 8 | 137 ± 3 | 107 ± 2 | 122 ± 4 |
| 20 | 63 ± 7 | 58 ± 1 | 51 ± 2 | 57 ± 4 |
| 21 | 48 ± 6 | 86 ± 10 | 56 ± 2 | 104 ± 2 |
| 22 | 10.1 ± 0.1 | 10 ± 3 | 19 ± 1 | 7.0 ± 0.3 |
| 23 | 75 ± 1 | 60 ± 7 | 119 ± 14 | 57 ± 5 |
| 24 | 42 ± 1 | 17 ± 2 | 81 ± 1 | 13 ± 1 |
| 25 | 57 ± 3 | 85 ± 5 | 17 ± 1 | 13.3 ± 0.2 |
| 26 | 8.3 ± 1.5 | 8.7 ± 1.9 | 13.4 ± 0.9 | 1.9 ± 0.3 |

<sup>a</sup> Extreme values – more than 100% or less than 10% – are given in bold. Values corresponding to low-, medium- and high activity are designated by blue, black and red, respectively (color table online). Ligand concentration 1 μM.

Unlike halogenation, introduction of amino group into position 2 (**13**) did not retain or rise the activity of the compound: the activity became lower compared to the parent BA (**1**) in plants expressing AHK3 and especially AHK2. Similar tendency of activity decrease was evident for the pair 6-benzyloxypurine (**14**) and its 2-amino-derivative (**16**).

Replacement of *N*^6^ with oxygen or sulfur led to a strong decrease in cytokinin activity of BA derivatives **14** and **15**, with clear difference: oxygen replacement (**14**) retained moderate activity while sulfur replacement (**15**) rendered the derivative low-active. Amino group introduction at *C*2 position (**16** vs **14**) reduced cytokinin activity even further, up to a negligible level in WT and mutant lines. The increase of the linker length in 2-amino-6-(2-phenylethoxy/propoxy)-purines (**17**, **18**) kept the cytokinin activity at the background level.

However, the activity of BA analogues with variable linker length was roughly inversely correlated with the number of methylene residues in the linker. The most potent was 6-phenyladenine (**19**), followed by unmodified BA (**1**). Compounds **20** and **21**, having two and three methylene groups, respectively, were on the whole moderately active, while longer linker (**22**) or bulkier aromatic residue (**26**) resulted in nearly complete loss of the cytokinin activity.

Compounds **23**–**25** showed some traits of selectivity between receptors. Disubstituted 2-chloro-6-furfuryladenine (**23**) and particularly 2-chloro-6-(1-phenylethyl)-adenine (**24**), though racemic, were most active in plants harboring AHK3 receptor. By contrast, monosubstituted 6-benzoyladenine (**25**) exhibited the highest activity in plants expressing AHK2 receptor (Table 2).

Despite these peculiar traits, receptors generally demonstrated rather high level of similarity in their response to various BA analogues, revealed by correlation analysis (Fig. 1). The lowest amount of outliers was observed upon comparison of plant lines bearing AHK2 and CRE1/AHK4 receptors (Spearman correlation coefficient RS = 0.87). Two other pairs of the double mutants were less similar (RS were 0.74–0.77).

**Fig. 1.**
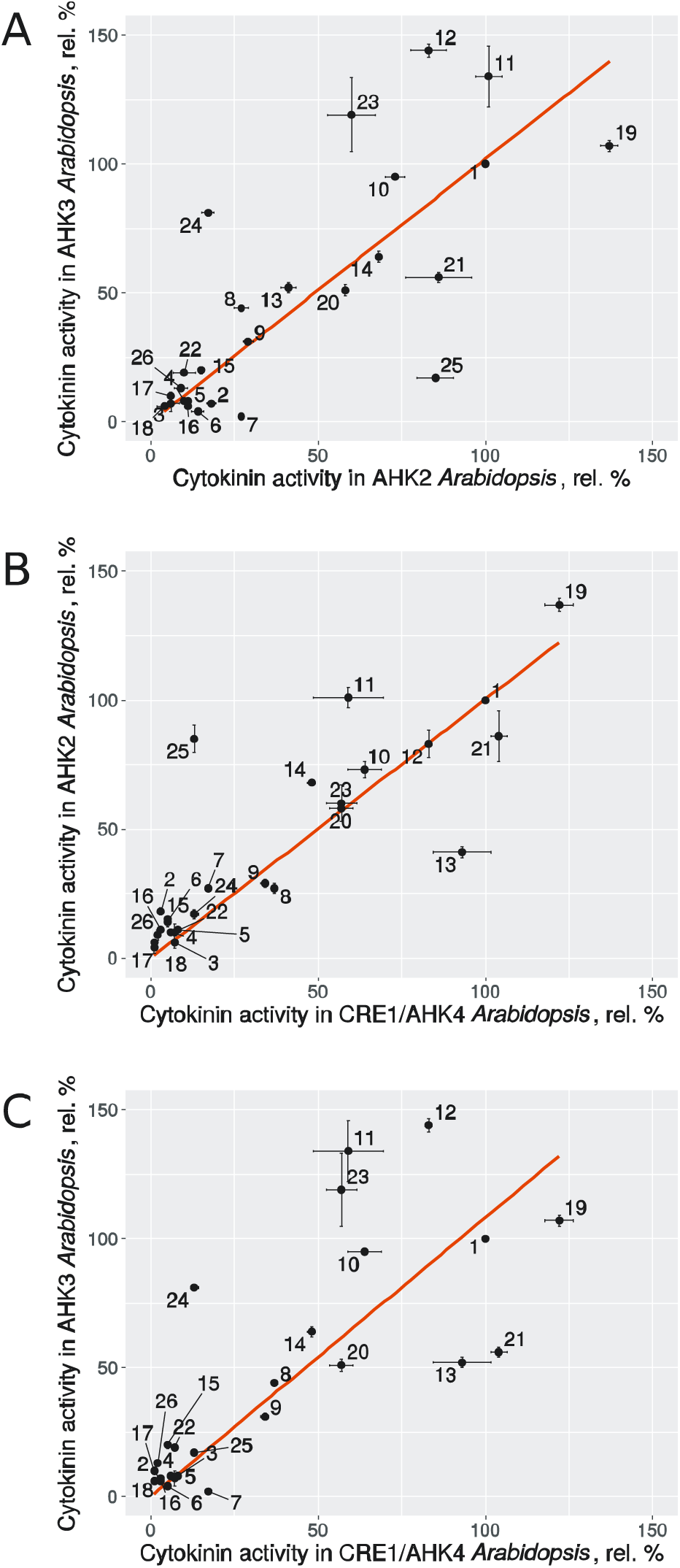
Correlations of cytokinin activity of tested compounds in *Arabidopsis* double mutants retaining only one isoform of cytokinin receptors (indicated as AHK*N Arabidopsis*). Axes show relative %. Ligand concentration in this and other assays was 1 μM.

### 3.2. *Cytokinin activity of BA analogues in the biotest with* Amaranthus *seedlings*

This assay is considered as a classical cytokinin biotest, rather specific and rapid. It is based on the cytokinin-dependent induction of a red pigment betacyanin synthesis in derooted *Amaranthus* seedlings in the darkness (Biddington and Thomas, 1973; Romanov et al., 2000). Most BA analogues screened for cytokinin activity in *Arabidopsis* double mutants were additionally tested using *Amaranthus* assay (Table 3). Both assays gave similar results for many BA derivatives, for example, no compound classified as high-active in one test system proved to have low activity in another one. Accordingly, BA derivatives halogenated at position 2 (**10**–**12**) usually had high cytokinin activity in both assay systems. Compounds bearing an amino group at *C*2 and *N*^6^ replacement by *O*^6^ (**16**, **17**) and especially compound **5** methylated at *N*9 were low-active in both biotests.

**Table 3.** Cytokinin activity of BA derivatives in biotest with *Amaranthus* seedlings ^a^.

| Compound | Relative activity (%) |
| --- | --- |
| **1** | **100** |
| **2** | 26 ± 5 |
| **5** | **7.6 ± 0.8** |
| **7** | 16 ± 5 |
| **9** | 24 ± 3 |
| **10** | 84 ± 8 |
| **11** | 100 ± 4 |
| **12** | 32 ± 5 |
| **13** | 85 ± 9 |
| **16** | 19 ± 5 |
| **17** | 18 ± 3 |
| **18** | 31 ± 1 |
| **19** | 78 ± 3 |
| **20** | 72 ± 4 |
| **21** | 81 ± 1 |
| **23** | 25 ± 3 |
| **24** | 26 ± 1 |
| **25** | 85 ± 5 |

<sup>a</sup> Extreme values – more than 100% or less than 10% – are given in bold. Values corresponding to low, medium and high activity are designated by blue, black and red, respectively (color table online).

However, clear differences between two assay systems were also observed. 9-Methyl-BA (**2**) bearing methyl group at position 9 was able to exert some cytokinin effect in *Amaranthus* but not *Arabidopsis* seedlings. BA derivatives **7**, **9**, **23** and **24**, classified as moderately active in *Arabidopsis* biotest, exhibited only low activity in *Amaranthus* seedlings. By contrast, compounds **13**, **21**, **25**, moderately active in *Arabidopsis* seedlings, had much higher cytokinin activity in *Amaranthus* assay (Table 3). Interestingly, in *Amaranthus* biotest the assayed compounds were divided into two groups, with relative activity between 7–32% or 72–100%; no intermediate (32–72%) activity was observed.

Despite aforementioned differences, results of the two assays were significantly correlated (Fig. 2). The best correlation with *Amaranthus*-derived data was observed for data obtained on either WT or AHK2 mutant *Arabidopsis* lines (RS = 0.68 and 0.72, respectively). In case of CRE1/AHK4 receptor, the correlation coefficient was slightly lower (RS = 0.59), and the worst correlation was observed in the case of AHK3 (RS = 0.55).

**Fig. 2.**
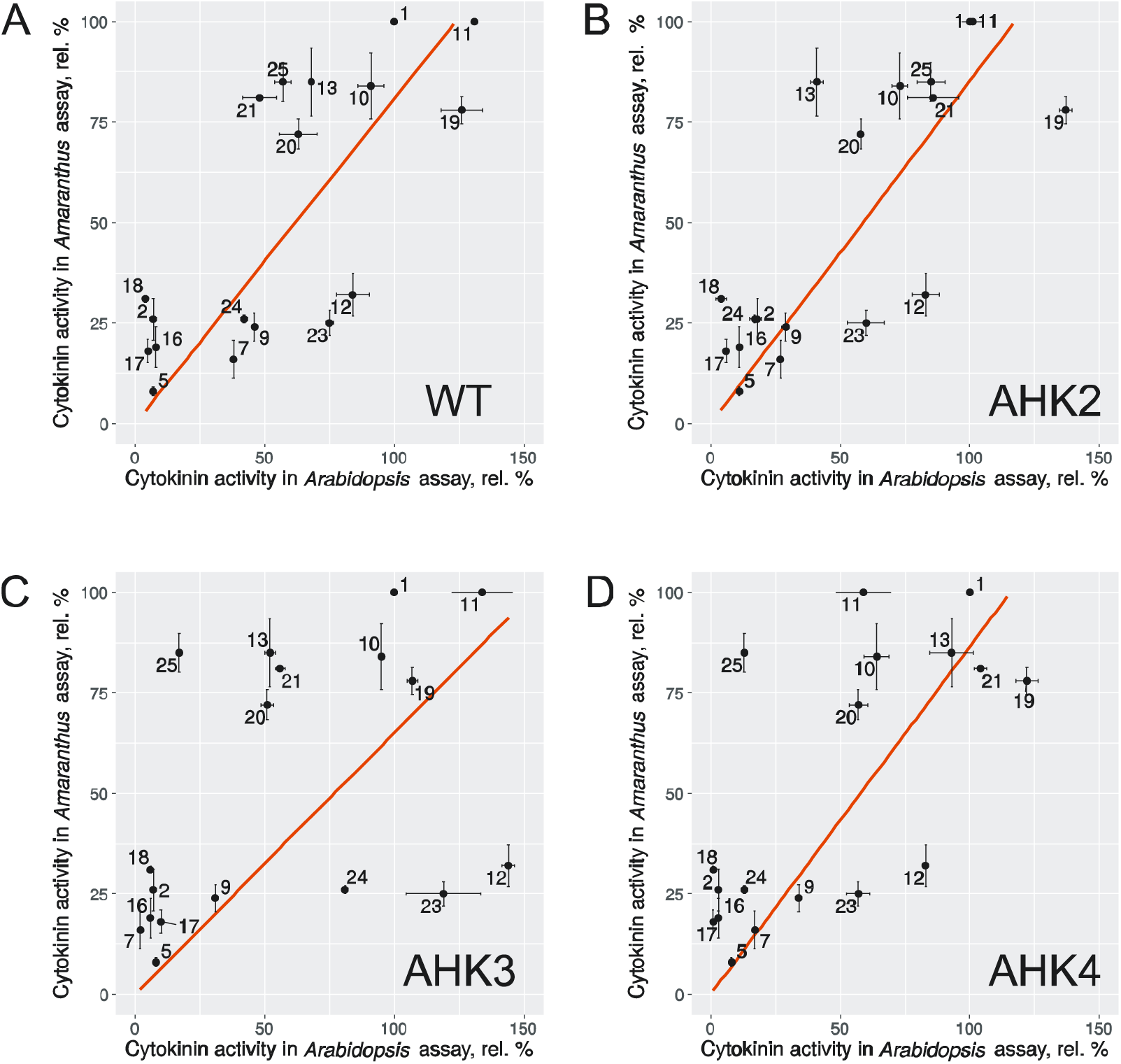
Correlations between cytokinin activity tested in *Arabidopsis* and *Amaranthus* seedlings. The plots show data on seedlings *Arabidopsis* vs *Amaranthus* plants: WT (A); expressing single receptor: AHK2 (B), AHK3 (C), or CRE1/AHK4 (D). Axes show relative %.

### 3.3. *Affinity of BA derivatives to individual cytokinin receptors from* Arabidopsis

Ligand recognition and receptor activation are grounded on high-affinity ligand-receptor interaction. We assessed the affinity of BA derivatives to each of *Arabidopsis* cytokinin receptors by means of two variations of the radioligand competition assay (Table 4). For AHK3 and CRE1/AHK4, a live-cell hormone-binding assay with transgenic bacteria expressing individual AHK receptors (Romanov et al., 2005; Romanov and Lomin, 2009) was employed. To provide better access of tested ligands to the expressed receptors, spheroplasts were obtained by removal of the bacterial shell (Romanov et al., 2005; Lomin et al., 2011). For AHK2, which was poorly cloned in *E. coli*, the plant assay system based on plant microsomes carrying transiently expressed receptors (Lomin et al., 2015) was used.

**Table 4.** Binding activity of BA-derivatives to individual cytokinin receptors of *Arabidopsis*^a^.

| Compound | Relative binding (%) to receptor: |  |  |
| --- | --- | --- | --- |
|  | AHK2 | AHK3 | CRE1/AHK4 |
| **1** | 93 ± 5 | 100 ± 6 | 88 ± 6 |
| **5** | 4.9 ± 0.2 | 50 ± 3 | 12 ± 1 |
| **7** | 14.5 ± 0.3 | 5.7 ± 0.7 | 27 ± 7 |
| **9** | 29 ± 3 | 28 ± 1 | 81 ± 3 |
| **10** | 99 ± 8 | 86 ± 3 | 89 ± 3 |
| **11** | 99 ± 6 | 100 ± 10 | 80 ± 5 |
| **12** | 92 ± 11 | 84 ± 4 | 86 ± 7 |
| **13** | 46 ± 2 | 77 ± 6 | 67 ± 6 |
| **14** | 52 ± 3 | 88 ± 6 | 91 ± 2 |
| **16** | 20 ± 1 | 0.7 ± 0.1 | 0.6 ± 0.1 |
| **17** | 14 ± 1 | 3.1 ± 0.3 | 7.9 ± 0.6 |
| **19** | 97 ± 3 | 66 ± 4 | 80 ± 5 |
| **20** | 89 ± 3 | 51 ± 5 | 48 ± 3 |
| **21** | 80 ± 3 | 90 ± 5 | 42 ± 2 |
| **24** | 30 ± 1 | 66 ± 5 | 8.9 ± 0.7 |
| **25** | 56 ± 2 | 2.2 ± 0.1 | 20 ± 2 |

<sup>a</sup> Extreme values – more than 100% or less than 10% – are given in bold. Values corresponding to low-, medium-, and high activity are designated by blue, black and red, respectively (color table online).

Several BA analogues were found to efficiently bind to every individual cytokinin receptor. This group included monosubstituted 2-halogenated BA derivatives (**10**, **11**), disubstituted 2-chloro-6-(3-fluorobenzyl)-adenine (**12**), and 6-phenyladenine (**19**). These compounds were usually highly active in both biotests (Tables 2, 3). Another group of BA analogues, which, on the contrary, did not almost bind to any of individual receptors, was represented by compounds **7**, **16**, **17**. All compounds of this second group exhibited small or almost no activity in both *in planta* assays (Table 2, 3). The only exception was 8-thio-BA (**7**), which showed medium activity in WT *Arabidopsis*. Compound **5**, which did not show cytokinin activity in both *in planta* assays, had expectedly low affinity to AHK2 and CRE1/AHK4, but surprisingly appreciable binding to AHK3 receptor. Other compounds were receptor-specific to a various extent. Among them were **9**, which preferentially bound to CRE1/AHK4; racemate 2-chloro-6-(1-phenylethyl)-adenine (**24**), which clearly preferred AHK3, and 6-benzoyladenine (**25**), which preferentially bound to AHK2.

The obtained data sets allowed to analyze the dependence between the cytokinin receptor activation by a compound and its binding affinity (Fig. 3). For all three receptors the correlation between GUS activity and binding level of the compound to the receptor was rather high (RS between 0.8–0.9) and the trend line started from axes zero interception. This underlines the importance of high-affinity binding of a ligand to the receptor for its recognition as a cytokinin signaling molecule. The best correlation (RS = 0.86–0.88) was observed for AHK2 and CRE1/AHK4, whereas in case of AHK3 (RS = 0.80) there were some few outliers.

**Fig. 3.**
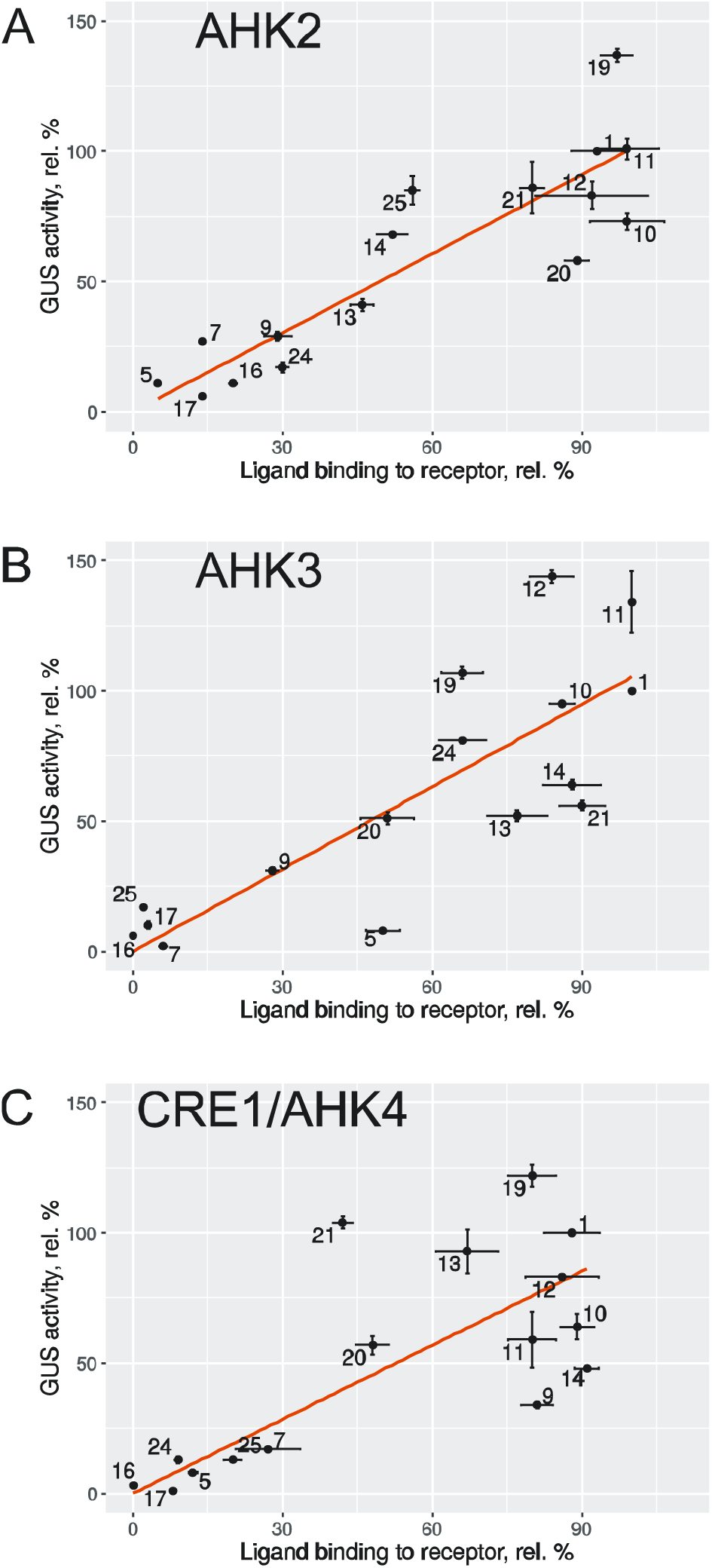
Correlations between ligand binding and hormonal activities of individual *Arabidopsis* receptors. Receptors: AHK2 (A); AHK3 (B); CRE1/AHK4 (C). Axes show relative %.

### 3.4. Molecular modeling of interaction between BA derivatives and cytokinin receptors

In order to rationalize structure factors underlying the binding efficiency of cytokinin analogues, we performed docking studies for them and optimized binding poses to obtain more meaningful estimates of interaction energy. Homology models of AHK2 and AHK3 were constructed using the CRE1/AHK4 X-ray structure complexed with *trans*-zeatin (PDB ID 3T4L) as the template (Lomin et al., 2015). Given very close similarity between X-ray structures of CRE1/AHK4, the most complete of them was chosen as the template. Three water molecules present in the cytokinin binding site of CRE1/AHK4 may play a crucial role in binding pose realization, prediction, and assessment. We analyzed two opposite configurations of the binding site: (1) configuration with all three waters present, denoted further with 'W', and (2) configuration with all three waters removed, denoted 'D' (from 'dry'). Water molecules were treated as rigid during the homology modeling and docking, and free movement was allowed to them during optimization.

Two different docking methods were employed for ligand pose generation for CRE1/AHK4 co-crystallized with ligands: FRED (McGann, 2011) takes the pool of ligand conformations generated by OMEGA and selects the best ligand orientation guided by the interactions with the binding site, whereas HYBRID (McGann, 2012) selects the best pose taking into account also the shape of the co-crystallized ligand. Best-scored solutions from FRED and HYBRID according to ChemGauss4 scoring function were optimized with SZYBKI after stripping or retaining waters. Because use of sole 3T4L structure as the model of CRE1/AHK4 could introduce bias, structures complexed with thidiazuron (3T4T) and *N*^6^-benzyladenine (3T4K) were also utilized in the same docking/optimization pipeline. Only homology models were available for AHK2 and AHK3, so HYBRID simulation for them was not performed.

Fig. 4 shows correlation between docking scores (from both FRED and HYBRID in 3T4L) and specific binding (SB) % for CRE1/AHK4. General expected trend for this correlation is highlighted by orange line: better binders should be scored better, obtaining more negative score values. Scores for poses generated by FRED and HYBRID were usually close to each other, with the only exception of 8-oxo-9-methyl-BA (**5**). Majority of compounds fell into quadrants I and IV, thus supporting the expectations. Two groups of outliers are present: good binder **9** (group 1) was scored similarly to poor binders **5**, **7** and **26**; poor binders **16**, **17**, **24** (both enantiomers) were scored similarly to other good binders.

**Fig. 4.**
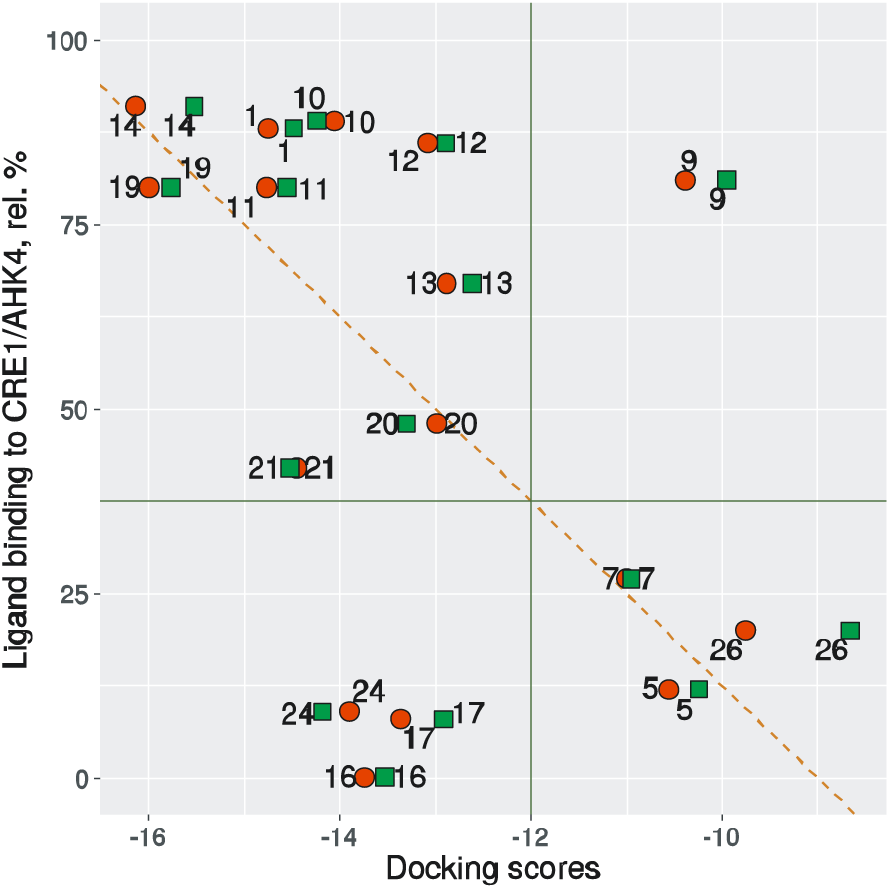
CRE1/AHK4 specific binding, relative %, vs 3T4L docking scores. Orange line: general expected trend. Quarter I (left top): good binders scored good. Quarter II (right top): good binders scored bad. Quarter III (left bottom): bad binders scored good. Quarter IV (right bottom): bad binders scored bad.

The most consistent overestimate was obtained for IE and docking scores of 2-amino-*O*^6^ analogues **16** and **17**. These compounds were predicted to bind very good, but experimentally observed binding was on a very low level. To rationalize these problems, we noticed that compound **16** bears two modifications compared with BA: amino group is introduced into position 2 and NH is substituted for *O* in position 6. Importantly, compounds with only one of these substitutions were already present in the set: **13** and **14**, respectively. Both of them bound to CRE1/AHK4 very efficiently, and predicted interaction energy and scores for them were overall favorable. All compounds from this series showed uniform binding modes (Fig. 5). Amino group in position 2 points to the space between a water molecule and a hydrophobic cavity, so unfavorable interactions are limited in this region. Oxygen atom in position 6, though, changes the character of possible interactions: it is a hydrogen bond acceptor, whereas NH is a hydrogen bond donor interacting with Asp262 of CRE1/AHK4. Nevertheless, Asp262 may be protonated to get the hydrogen bond back; alternatively, removal of one hydrogen bond may not be crucial for overall binding efficiency. Thus, it is not unexpected that **16**, combining two tolerable substitutions, is scored as favorably as each of **13** and **14**. Compound **17** bears one more modification compared with **16** — benzyl moiety is substituted with 2-phenylethyl one. If such a substitution is performed on BA, compound **20** appears, and its specific binding percent is about half of BA's, but calculated interaction energy and scores are very favorable due to higher hydrophobic surface area and size well tolerated by the binding site. Applying the same reasoning that was used above for **16**, it is again not unexpected for **17** to receive favorable scores, but all these modifications combined in a single molecule are apparently not tolerated by the receptor. There were no 2-phenylethyl analogues of **13** and **14** in the set, but they are expected to bind poorer and to be less effective than parent compounds.

**Fig. 5.**
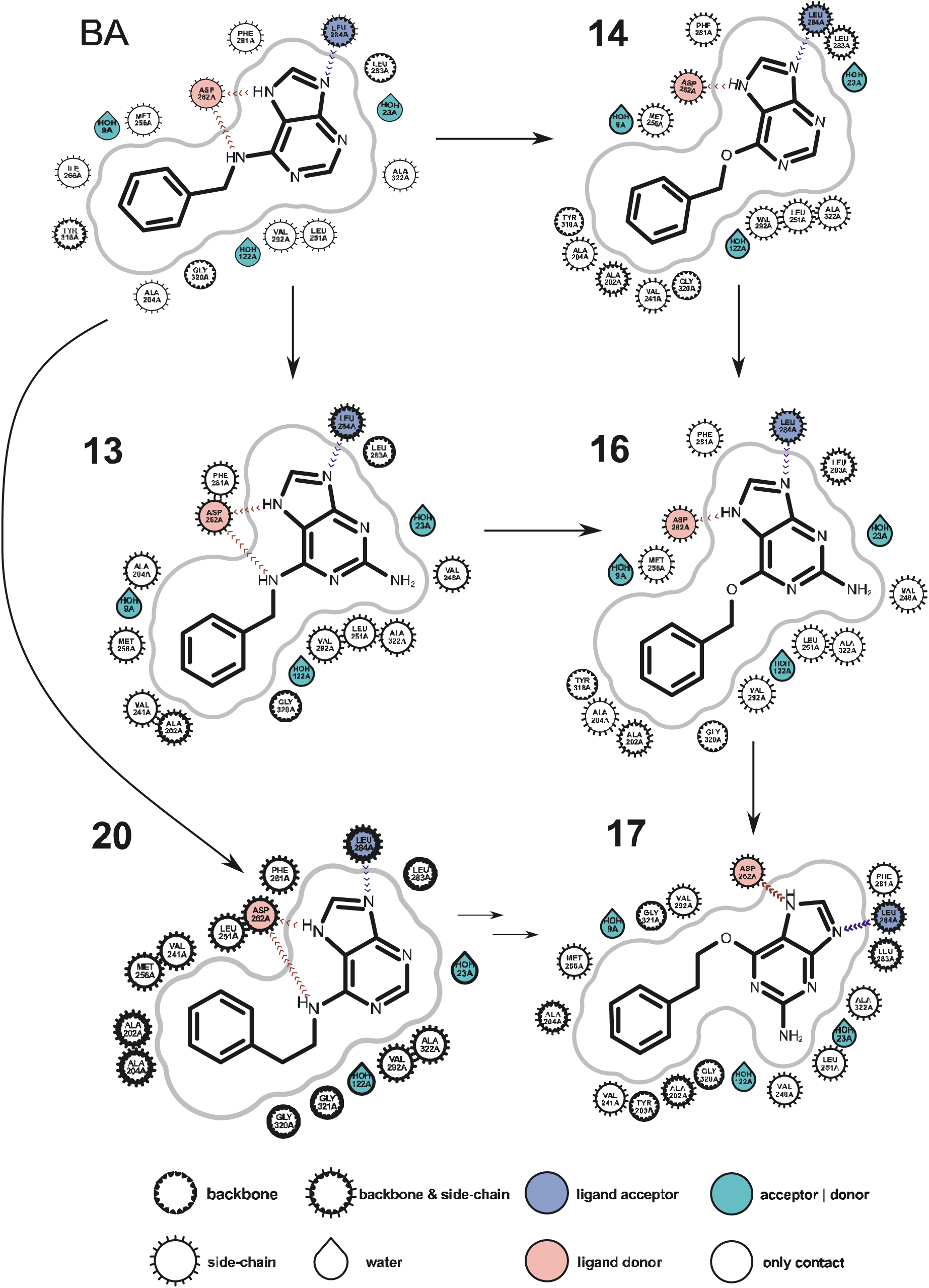
2D representations of 3T4L docking results of BA (**1**), **13**, **14**, **16**, **17**, **20**. Hydrogen bonds are shown as dotted lines.

It should be noted that the pose of BA as observed in the X-ray structure (3T4K), despite well optimized by the authors of the structure without bumps between the ligand and receptor, could not be reproduced perfectly by the docking procedure due to selection of a slightly different ligand conformation. In this conformation the benzyl group of BA contacts with Ala202 a bit too close, thus leading to unfavorable score contributions from two C atoms of benzyl (Fig. 6). Orientation of benzyl in **13** and **14** complies with the binding site better, but unfavorable contributions are received from oxygen atom and NH^2^ group, respectively (Fig. 7). Overall scores for BA, **13** and **14** are thus similar. More pronounced bumps are observed for the docked poses of **17** and **20**, where 2-phenylethyl needs to be tightly packed in the binding site. Although there is still enough space to accommodate longer side chains (compare **21** with three methylene groups between *N*^6^ and phenyl), packing becomes tighter, and selectivity between the receptors may arise due to differences in the amino acid composition of the binding site.

**Fig. 6.**
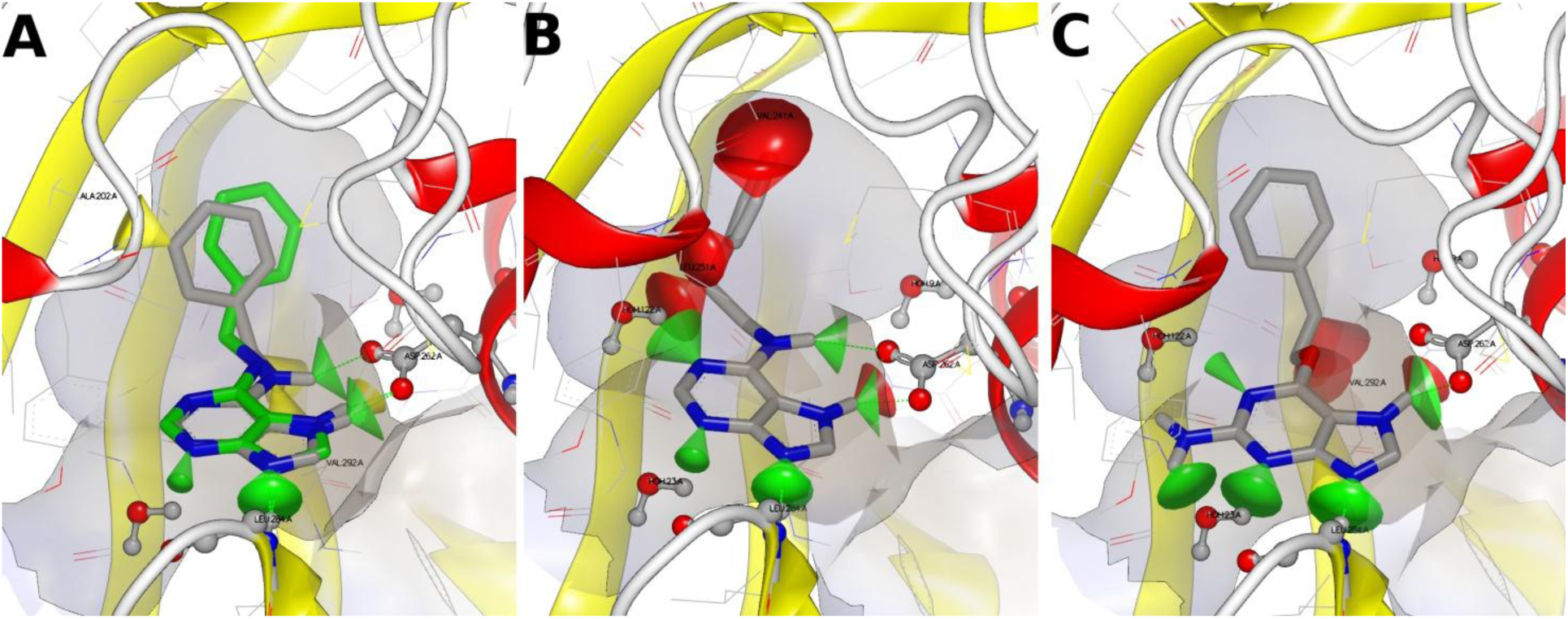
Binding modes of BA, **20**, and **17** according to docking. A) Docked pose of BA (**1**, gray carbons) superimposed with X-ray pose of BA (**1**) in 3T4K (green carbons). Note the yellow cone of less favorable interaction with Ala202 for the docked pose. B, C) Docked poses of **20** (B), and **17** (C) into 3T4L. Cones denote the interactions: green, favorable; yellow, less favorable; red, unfavorable.

**Fig. 7.**
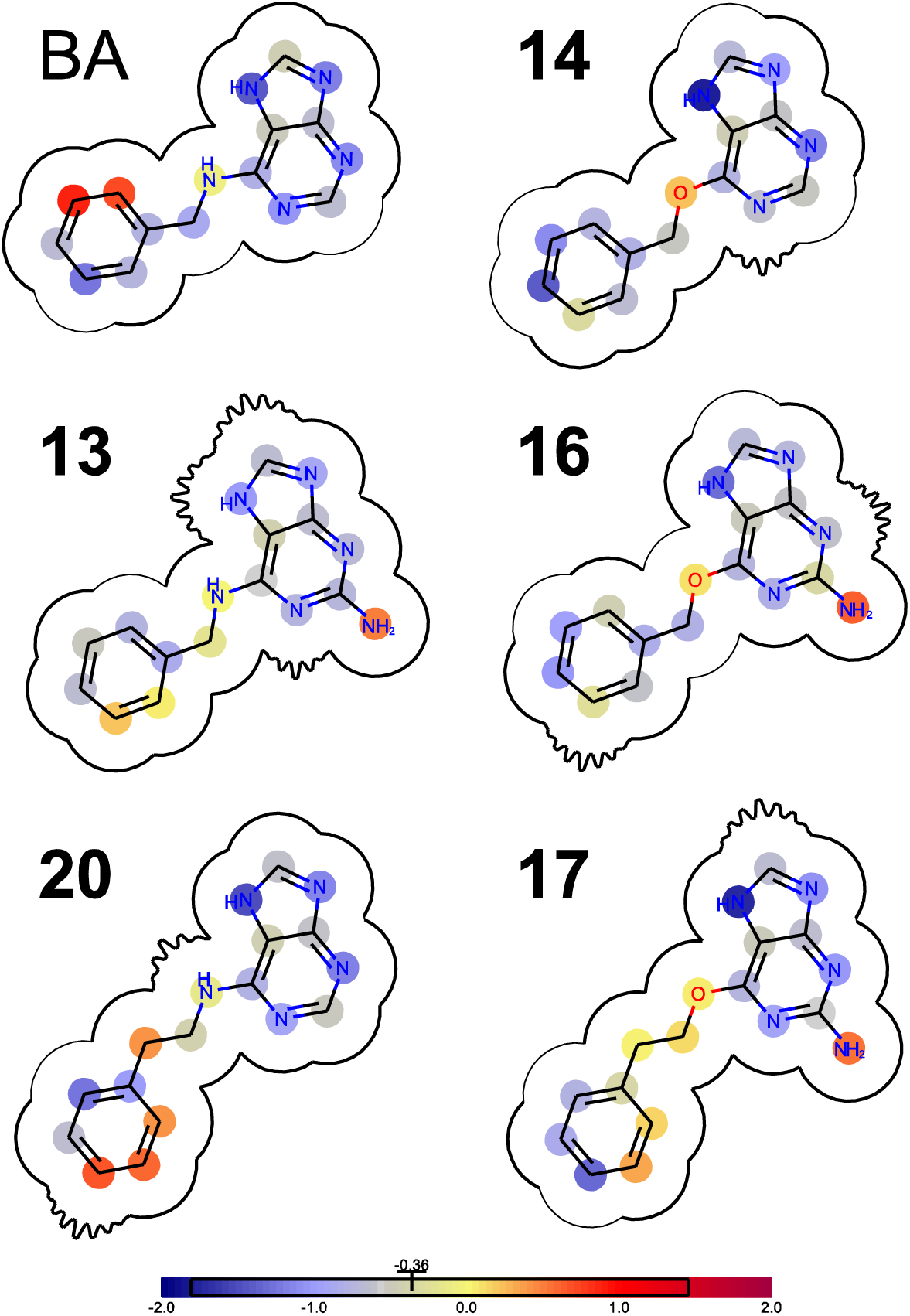
2D representations of 3T4L docking results for BA (**1**), **13**, **14**, **16**, **17**, **20**. Protein cavities are shown with kinked curves; atoms are highlighted according to score contributions. Color scheme for score contributions in ChemGauss4 units is in the bottom.

Another position of adenine ring that significantly affects ligand binding properties is position 9. The active tautomer of adenine in cytokinins is protonated at *N*7, and *N*9 is not protonated, thus accepting a hydrogen bond from Leu284 NH. Change of hydrogen bond donor/acceptor pattern at *N*7 and *N*9 leads to changes in ligand binding modes. One option for such a change is methylation at *N*9, realized in compounds **2–5**, prohibiting protonation of a neutral cytokinin molecule at *N*7 (Fig. 8). Despite ligand can still enter the binding site, the perfect hydrogen bonding pattern cannot be realized. Reorientation of the ligand makes binding technically possible, but receptor activation with these molecules is not efficient.

**Fig. 8.**
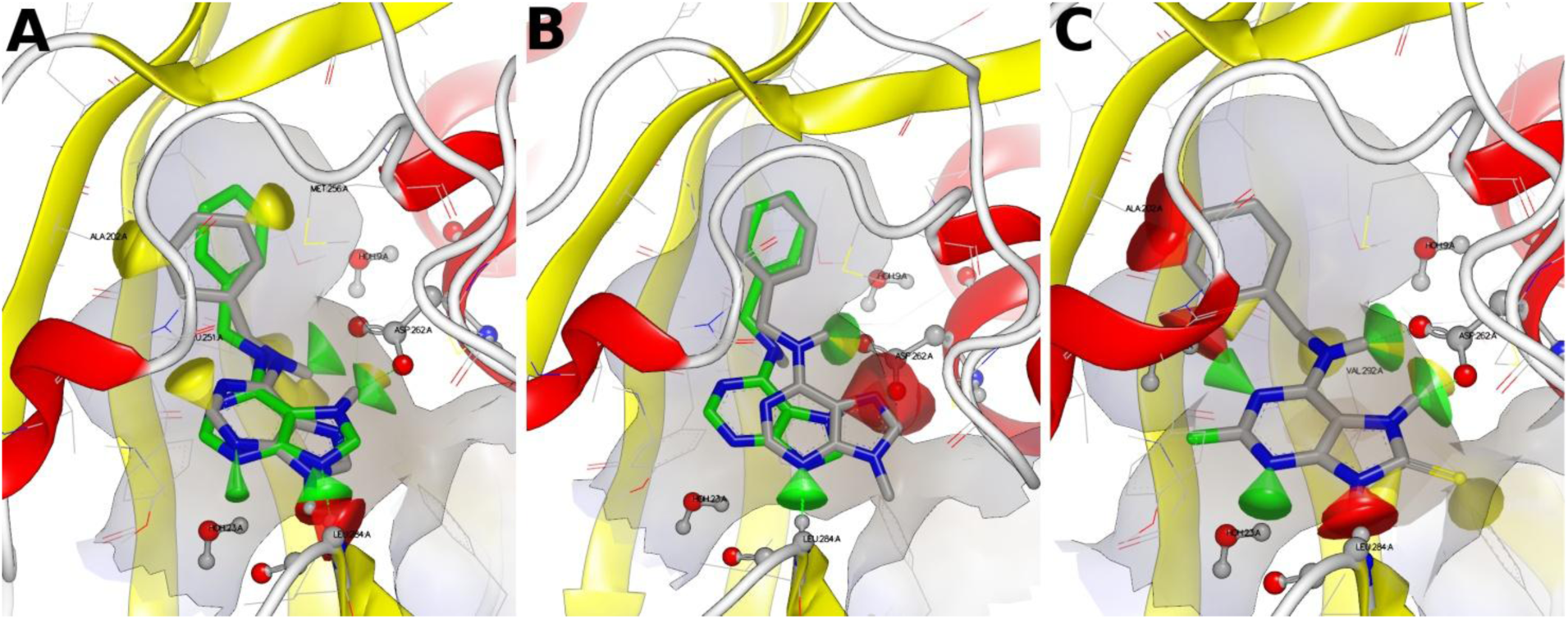
Binding analysis of 9-substituted cytokinin analogues. A) Superimposition of 9-methyl-BA (**2**, gray) with X-ray orientation of BA (green) in the ligand binding site of 3T4K. Note a significant bump between the methyl group and Leu284. B) Docked pose of **2** (gray) superimposed with X-ray orientation of BA (green). Hydrogen bond with Leu284 is realized through *N*3, but *N*7 unfavorably interacts with Asp262. C) Docked pose of **9**. Despite hydrogen atom at *N*9 unfavorably interacts with Leu284, hydrogen bonds with Asp262 are efficient enough.

Position 9 of the adenine moiety may also be protonated along with position 7, as exemplified by compounds **6**–**9**. According to quantum chemical calculations (Supplementary Table 2), tautomeric equilibrium of these compounds is completely shifted to the tautomer B (Table 1; Supplementary Table 1), with hydrogen atoms at both *N*7 and *N*9. Hydrogen bond donor at *N*9 cannot form a hydrogen bond with backbone NH of Leu282, which is present in all the crystal structures; nevertheless, significant levels of affinity and activity for **9** show that imperfect interaction does not guarantee absence of activity. Ligand molecule can also be re-oriented in the binding site of CRE1/AHK4 to form a hydrogen bond between *N*3 and Leu282 NH. Another option is change of Leu284 loop orientation, which is not sampled in the docking to the rigid protein. During SZYBKI optimization re-orientation of **9** is sufficient for achieving a favorable interaction energy.

Compound **25** is not common for this series due to presence of an internal hydrogen bond between the carboxyl oxygen and *N*7 hydrogen (Fig. 9). This interaction is very favorable for the ligand itself, but this ligand may form one hydrogen bond less with the protein. Consequently, adenine moiety is predicted to be inverted in the binding site, and N3 appears to accept hydrogen bond from Leu284 NH again. This lack of ligand-receptor hydrogen bond nicely explains low affinity and activity of this compound against AHK3 and CRE1/AHK4. Surprisingly, in the model of **25** — AHK2 complex more tight binding with five hydrogen bonds is realized, ensuring higher affinity.

**Fig. 9.**
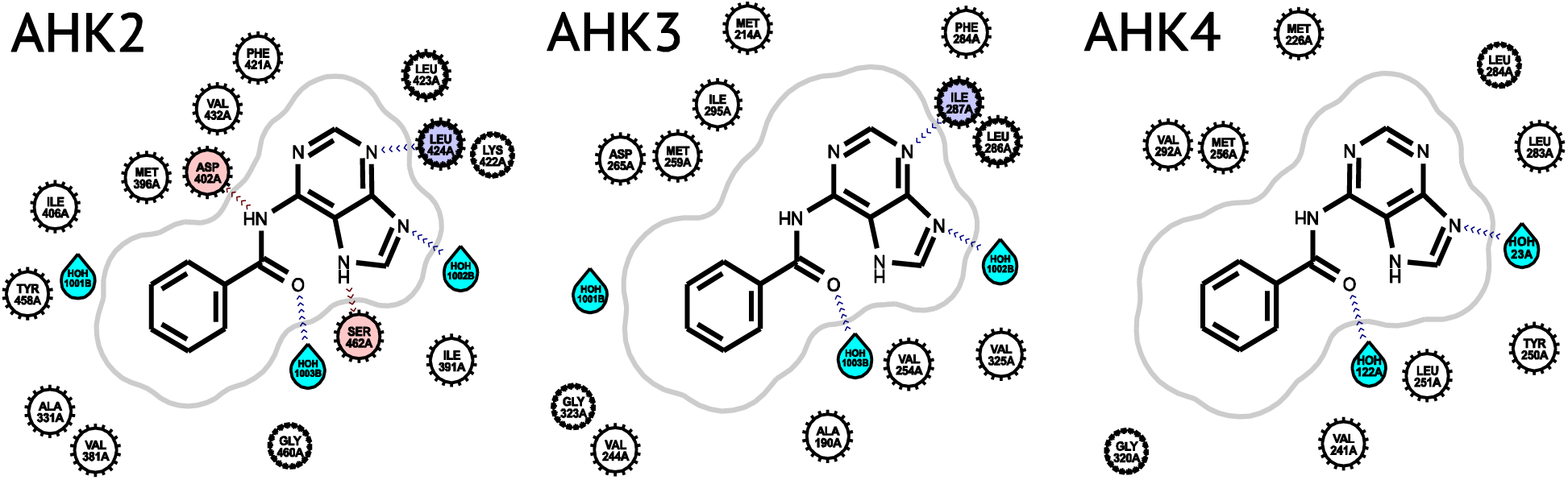
Predicted binding mode of **25** to each of cytokinin receptors of *Arabidopsis*. Adenine moiety is inverted with regard to BA.

Despite correlation between cytokinin activity and specific binding for AHK2 is the highest among the three receptors studied here, correlation of the docking scores (Fig. 10A) or interaction energy values (Fig. 10C) with specific binding % are much worse. Outliers are numerous for docking results, showing that the rigid model does not adequately represent the binding site. After optimization trends in the data are more prominent, but still the same compounds **13**, **16**, **17**, and **24** are scored too high. Presence of water dramatically affects interaction energy values for **14** and **25**, confirming limited reliability of the model. Nevertheless, remaining compounds group in the expected regions of the plot.

**Fig. 10.**
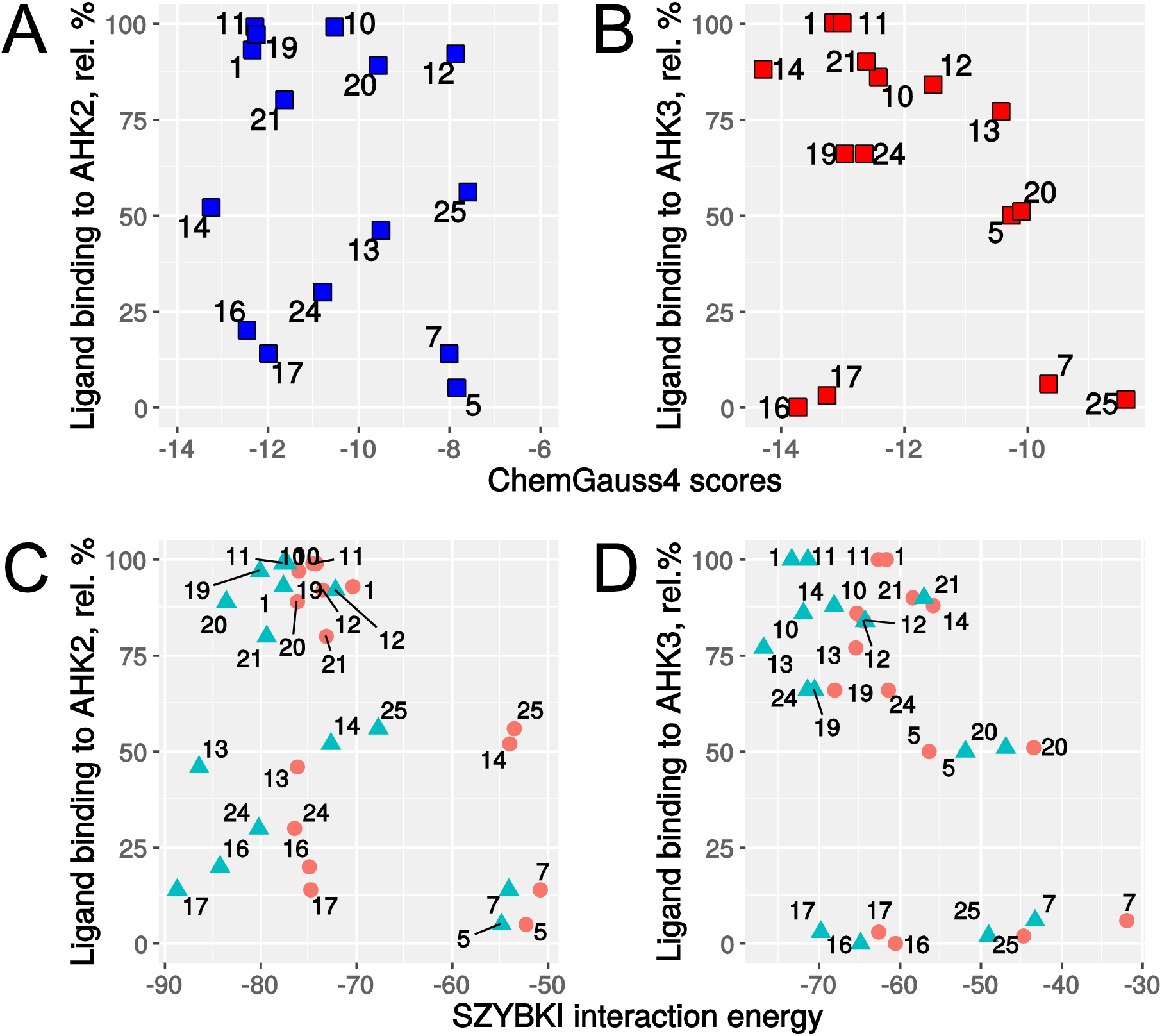
Correlation of AHK2 and AHK3 specific binding with docking scores and SZYBKI interaction energy. A) AHK2 specific binding (relative %) versus ChemGauss4 scores from FRED docking. B) AHK3 specific binding (relative %) versus ChemGauss4 scores from FRED docking. C) AHK2 specific binding (relative %) versus SZYBKI interaction energy. Interaction energy values for **9** (no water: +54.74; with water: +56.65) are far outside the typical range. D) AHK3 specific binding (rel. %) versus SZYBKI interaction energy.

The most prominent correlation between docking scores and specific binding % is observed for AHK3. All the points are lined along the general expected trend line, with the only exceptions of **16** and **17**, which are scored reasonably good again (Fig. 10B), but show weak binding (Table 4). Situation does not become any better with SZYBKI optimization (Fig. 10D), moreover, interaction energy of the optimized pose of **9** is highly unfavorable up to the positive values. No other outliers appeared.

## 4. Discussion

Cytokinin receptors constitute in plants small families of a few isoforms, differing in their ligand specificity and expression pattern (Stolz et al., 2011; Lomin et al., 2012; 2015). To study characteristics of individual cytokinin receptors, double receptor mutants of *Arabidopsis*, each expressing a different single receptor, were employed (Riefler et al., 2006). These mutants were supplied with cytokinin-sensitive *P_ARR_*_5_*:GUS* construct allowing quantitative measurements of cytokinin effects (Stolz et al., 2011). Importantly, this assay system is intended to analyze and compare properties of individual receptors functioning *in planta*. Other advantages of the system are its simplicity, rapidity, and quantitative character. To our knowledge, *P_ARR_*_5_*:GUS-*equipped double receptor mutants were employed in this work for massive cytokinin activity screening for the first time.

By means of this assay system, we have screened 25 BA derivatives for cytokinin activity. The hormonal potential of these substances was further studied in the traditional assay with derooted *Amaranthus* seedlings. For many compounds, there was a marked correlation between two biotests, the ligand specificity of *Amaranthus* receptors resembled that of AHK2 rather than AHK3. The obtained results were also in general agreement with previous data from different assays where some substances of interest were investigated (Matsubara, 1990). BA modifications usually led to the reduction of the cytokinin activity. The rare exception was a halogen (F, Cl) substitution at *C*2 of the purine moiety (**10**, **11**). Such substitution can even partially restore the activity of BA derivatives severely inactivated by switch to scaffold B (**7**–**9**). Assays on double receptor mutants has shown that these halogenic substituents increase the compound activity preferentially towards the receptor AHK3 (**8**–**12**, **23**, **24**). On the contrary, amine residue at this position markedly reduced the overall cytokinin activity and particularly that towards the AHK2 receptor (**13**). In agreement with earlier data (Matsubara, 1990), the substitution of *N*^6^ by oxygen (*O*^6^) and especially by sulphur (*S*^6^) significantly decreased the cytokinin activity of the compounds (**14**, **15**) for all AHK receptors. A combination of two substitutions, namely *O*^6^ instead of *N*^6^ and amine group at *C*2, further reduced the activity of the compounds (**16**–**18**) up to ca. background level.

A comparison of ligand preferences of *Arabidopsis* receptors showed an overall correlation between receptors (Fig. 1), corroborating a high degree of similarity of their cytokinin binding sites (Steklov et al., 2013). Among all three receptor pairs, AHK2–CRE1/AHK4 one was distinguished by the highest similarity of activity profiles. This result is in accordance with previous observations on natural cytokinins (Stolz et al., 2011; Lomin et al., 2012). To note, the extent of divergence of hormone-binding CHASE domains is rather similar for all receptor pairs, homology degree is each case is around 63–64%.

In our study, we have supplemented data on cytokinin activity *in planta* of numerous BA derivatives with data on efficiency of ligand binding to the same individual cytokinin receptors. Binding assays were performed by two *in vitro* methods: live-cell assays using *E. coli* spheroplasts transformed with genes for individual receptors AHK3 or CRE1/AHK4 (Romanov et al., 2005; Lomin et al., 2011), and assays with microsomes isolated from tobacco leaves transiently expressing AHK2 receptor gene (Lomin et al., 2015). Recent studies showed that these two methods are consistent and equally valid when analyzing cytokinin-like ligands in base form (Lomin et al., 2015). Therefore data obtained by one of the methods compare well with analogous data obtained by another method.

The use of different approaches allowed us to compare the ability of a ligand to bind to the receptor with the ability of the same ligand to trigger signaling of the same receptor *in planta*. It is commonly accepted that a high-affinity binding of a ligand to a receptor is a prerequisite for receptor activation and signaling. Our data are in accordance with this postulate as regards *Arabidopsis* cytokinin receptors: the Spearman coefficients of correlation between binding and activation were as high as 0.80–0.88. However, this tight correlation does not preclude from the occurrence as outliers of rare cytokinin analogues exhibiting anticytokinin activity. Such antihormones strongly bind to receptors but do not activate them, thus competing with genuine hormones for receptor binding sites. Some synthetic receptor antagonists representing distinct BA derivatives were discovered some years ago (Spíchal et al., 2009; Nisler et al., 2010; Krivosheev et al., 2012). Among these novel anticytokinins, 6-(2-hydroxy-3-methylbenzylamino)purine (PI-55) (Spíchal et al., 2009) and 6-(benzyloxymethyl)adenosine (BOMA) (Krivosheev et al., 2012) were found to be receptor-specific, blocking CRE1/AHK4 rather than AHK3.

Receptor-specific compounds are of particular interest since they represent valuable tools to regulate the activity of individual receptors *in vivo*. In the present study new receptor-specific regulators were found. Compounds **24** and **25** behaved as specific agonists for the receptors AHK3 and AHK2, respectively. Data on preferential receptor activation (Table 2) were supported by data on preferential binding (Table 4). Notably, both compounds bear substituents in the linker between purine and aromatic parts of the molecule. This highlights the role of the linker modifications in rendering BA derivatives receptor-specific. Additional substituent in **24**, namely chlorine at position 2 of the purine heterocycle, confirms our hypothesis that AHK3 prefers BA derivatives with a halogen at this position. It should be noted that compound **24** has a chiral centre at the linker carbon atom and thus exists as two enantiomers. In the present study a racemic mixture of enantiomers was employed, so the exact role of each enantiomer remains to be elucidated. In addition, compound **5** has drawn our attention by its ability to bind preferentially to the AHK3 receptor without its activation, thus exhibiting some features of anticytokinin, AHK3 receptor specific antagonist. All these newly revealed receptor-specific compounds are promising for further studies.

To better clarify the observed features of ligand-receptor interaction, protein structure analysis and molecular modeling were employed. The scheme shows hydrogen bonding pattern between BA and CRE1/AHK4 according to X-ray structure (PDB ID 3T4K) (Hothorn et al., 2011). The *N*9 atom forms a hydrogen bond with the nitrogen atom of the main chain of Leu284, which is always protonated. Hence, BA molecule binds in the form where the *N*7 atom must be protonated. This last condition cannot be fulfilled in case of *N*9-substitutions which explains why *N*9 alkylated derivatives **2**–**5** are poor cytokinins. Based on the structure of CRE1/AHK4–BA complex, it was estimated that the introduction of bulky substituents at positions 2 or 8 of BA causes steric hindrance due to unfavorable contacts (less than 3 Å) of the substituent atoms with the side chain atoms of Val248 and Phe281 residues, respectively. However, tested BA derivatives had not enough bulky substituents at indicated positions, so the observed decrease in cytokinin activity of corresponding ligands was due to other reasons.

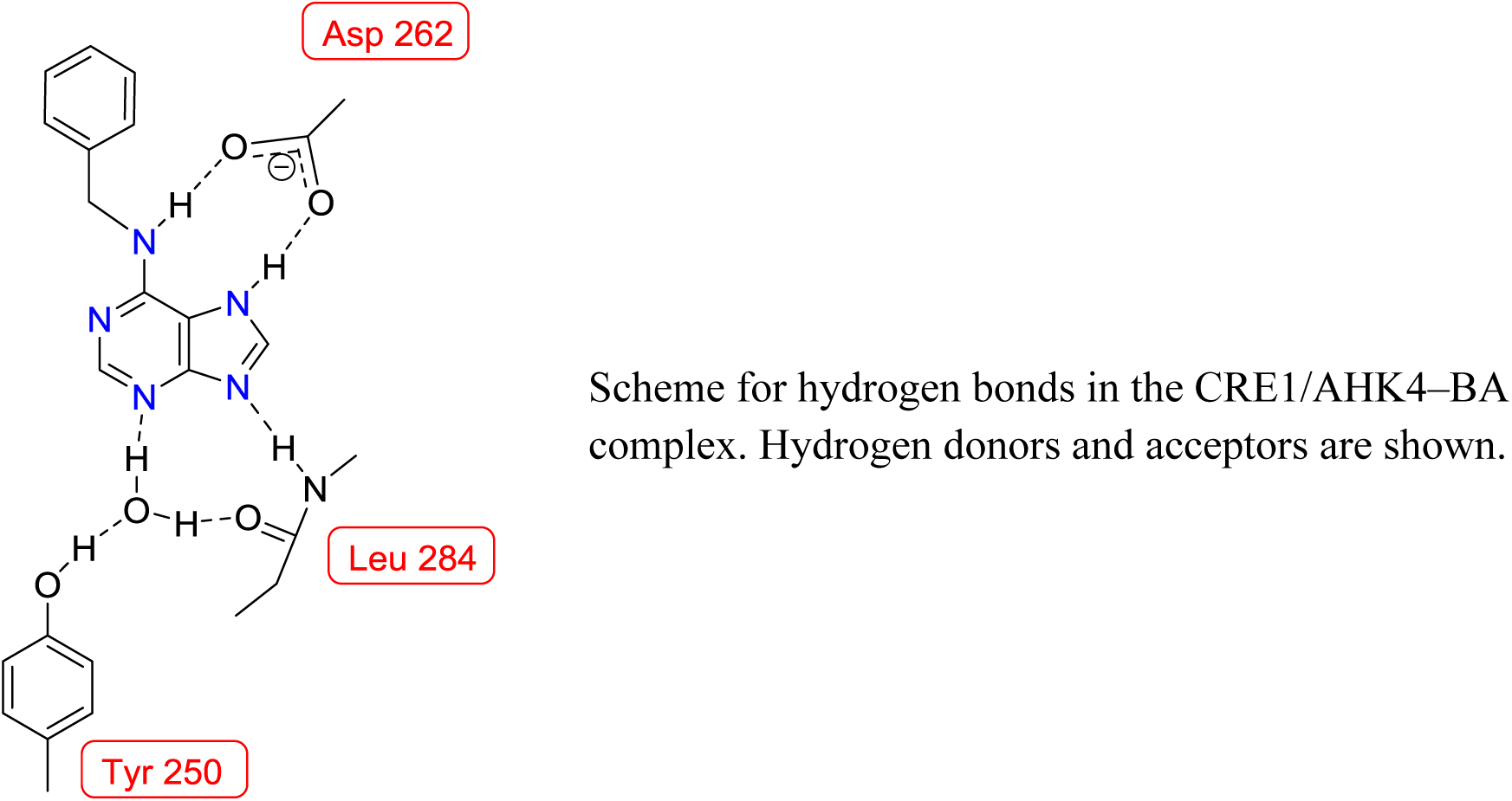
Scheme for hydrogen bonds in the CRE1/AHK4–BA complex. Hydrogen donors and acceptors are shown.

By the use of molecular modeling and docking, possible explanations for the observed differences in the ligand preferences of the receptors were obtained. For example, derivative **25** demonstrated clear preference towards receptor AHK2. This compound was predicted to bind to each receptor in a distinctive configuration, the adenine moiety being inverted with regard to BA.

According to our modeling, the AHK2 binding site was optimized so favorably that the hydrogen bond between Asp402 (homologue of Asp262) and NH of the ligand was retained. Moreover, when this ligand is complexed with AHK2, nearly all potential hydrogen bonds were realized after model optimization. In other pairs of the ligand-receptor, the crucial hydrogen bond was not conserved and the binding site was not filled. The modeling showed that the advantage of AHK2 may be due to the fact that its binding site is more compact than in other receptors where too few hydrogen bonds with **25** can be formed (Fig. 9).

An attempt to correlate experimental and modelling results was made (Figs 4, 10). Some of these plots, particularly those for AHK3, show reasonable correlations between experimental and theoretical data (Fig. 10). This result is promising for the rational design of new ligands with predictable cytokinin activity, particularly specific receptor agonists or antagonists.

## 5. Conclusion

The ligand preference of each of known cytokinin receptors of *Arabidopsis* (AHK2, AHK3, and CRE1/AHK4) was assayed *in planta*, by measuring the activity of cytokinin-sensitive promoter, and *in vitro*, by determining the ligand-receptor binding. Double receptor *Arabidopsis* mutants and 25 BA derivatives with various substituents in purine heterocycle, linker or aromatic moiety were used. Prominent correlations between ligand specificity profiles of different receptors corroborate the functional similarity of their hormone-binding sites, particularly those of AHK2 and CRE1/AHK4. Nevertheless, ligand binding properties of receptors are not identical, each individual receptor was shown to be distinctive in its ligand preference. As was expected, the ability of ligand to bind to the receptor was highly correlated with the intensity of the receptor signaling. Imperfect correlation between ligand binding and receptor activation underlines the complexity of agonist-triggered conformational rearrangements of the cytokinin receptors, when high affinity binding of a ligand is a necessary but not sufficient condition for "switching on" the receptor. Among tested BA derivatives, a few compounds were found which activate preferably one of receptors. These new receptor-specific cytokinins are promising for further studies. Receptor structure analysis, molecular modeling and molecular docking were employed to rationalize the interaction patterns between individual receptors and the ligands. The obtained dependencies between docking scores and experimental binding data provide a basis for design of new receptor-specific cytokinin analogues.

## 6. Experimental procedures

### 6.1. Chemicals

BA was used in the experiments as a standard. Ligand stock solutions in 100% DMSO at a concentration of 0.1 M were diluted to the desired concentrations with distilled water, DMSO concentration in 1 μM solutions was as small as 0.001%. Compounds were assayed at concentration 1 μM. For synthesis and properties of individual compounds, see *6.4*.

### 6.2. Cytokinin activity assays

Cytokinin activity of the compounds was evaluated in two assay systems based on *Arabidopsis thaliana* (L.) Heynh. and *Amaranthus caudatus* (L.) seedlings. In the first case, we used double mutants of *Arabidopsis*, in which only one of three cytokinin receptors (AHK2, AHK3 or CRE1/AHK4) was kept active (Riefler et al., 2006). All *Arabidopsis* plants used including WT control were transformed with the reporter *GUS* gene driven by cytokinin-dependent promoter PARR5 (Stolz et al., 2011). These plants were kindly provided by Prof. T. Schmülling and Dr. M. Riefler. 4–5 day-old *Arabidopsis* seedlings were incubated for 16 h in aqueous solutions of compounds (Romanov et al., 2002). Cytokinin activity of each compound was quantified by determination of the level of GUS activity reflecting the intensity of the *P_ARR_*_5_*:GUS* expression (Zvereva and Romanov, 2000). All 25 compounds (+ BA) shown in Table 1 and Supplementary Table 1 were tested in this assay system.

In the assay system based on *Amaranthus* seedlings, 17 of 25 compounds were tested. For that, 3–4 day-old etiolated and derooted *Amaranthus* seedlings were incubated for 16 h in the dark in solutions of compounds. Cytokinin activity of each compound was quantified by optical determination of the level of red pigment amaranthin accumulation in cotyledons (Biddington and Thomas, 1973; Romanov et al., 2000). In all our experiments, probes with BA and plain water were included as positive and negative controls, respectively. Cytokinin activity of compounds was evaluated as percentage relative to the BA activity at the same concentration in the same experiment.

### 6.3. Cytokinin binding assays

Direct binding of compounds to *Arabidopsis* cytokinin receptors was assessed by radioligand method. Experiments with AHK3 and CRE1/AHK4 receptors were performed in the assay system based on transformed *Escherichia coli* strain KMI001 (Romanov et al., 2005). *E. coli* were grown overnight at 25 °C in LB medium supplemented with 30 μg/ml chloramphenicol or 100 μg/ml ampicillin for bacteria expressing AHK3 or CRE1/AHK4 gene, respectively. All further operations were carried out at 4 °C. Spheroplasts of the grown bacteria were obtained as described (Lomin et al., 2011).

Assay system for AHK2 receptor was based on plant membranes isolated from transiently transformed leaves of *Nicotiana benthamiana*. Procedures for transient expression of AHK2 receptor gene in tobacco plants, plant membrane isolation and hormone binding assays were described in detail in (Lomin et al., 2015). Notable dissimilarity from the referred procedure was the use of tritium labeled isopentenyladenine (^3^H-iP, 2 pmol per probe, sp. act. 17.4 Ci/mmol) (Sidorov et al., 2015) instead of tritium-labeled *trans*-zeatin.

To estimate the binding of the ligand to the receptor, the amount of specific binding was determined as difference between total and nonspecific (with excess of unlabelled ligand) binding. Unlabelled ligands were used at concentration 1 μM. The level of specific binding of BA was taken for 100%. The binding level of a distinct ligand to the receptor was determined as % relative to specific BA binding.

### 6.4. Synthesis

#### General

The solvents and materials were reagent grade and were used without additional purification. Column chromatography was performed on silica gel (Kieselgel 60 Merck, 0.063–0.200 mm). TLC was performed on Alugram SIL G/UV254 (Macherey-Nagel) with UV visualization. Melting points were determined with Electrothermal Melting Point Apparatus IA6301 and are uncorrected. ^1^H and ^13^C (with complete proton decoupling) NMR spectra were recorded on Bruker AMX 400 NMR instrument at 303K. ^1^H-NMR-spectra were recorded at 400 MHz and ^13^C-NMR-spectra at 100 MHz. Chemical shifts in ppm were measured relative to the residual solvent signals as internal standards (CDCl_3_, ^1^H: 7.26 ppm, ^13^C: 77.1 ppm; DMSO-*d*_6_, ^1^H: 2.50 ppm, ^13^C: 39.5 ppm). Spin-spin coupling constants (*J*) are given in Hz. High resolution mass spectra (HRMS) were registered on a Bruker Daltonics micrOTOF-Q II instrument using electrospray ionization (ESI). The measurements were done in a positive ion mode. Interface capillary voltage: 4500 V; mass range from m/z 50 to 3000; external calibration (Electrospray Calibrant Solution, Fluka); nebulizer pressure: 0.4 Bar; flow rate: 3 µL/min; dry gas: nitrogen (4L/min); interface temp. 200 °C. Samples were injected in to the mass spectrometer chamber from the Agilent 1260 HPLC system equipped with Agilent Poroshell 120 EC-C18 (3.0 × 50 mm; 2.7 µm) column; flow rate 200 µL/min; samples were injected from the acetonitrile-water (1:1) solution and were eluted in a linear gradient of acetonitrile concentrations (50 → 100%).

The following 6-substituted purine derivatives were obtained according to the previously published protocols: 2-chloro-6-benzylaminopurine (**10**) (Theiler et al., 1976), 2-amino-6-benzylaminopurine (**13**) (Wang et al., 2013), 6-benzyloxypurine (**14**), 6-phenylaminopurine (**19**), 6-(2-phenylethyl)-aminopurine (**20**), 6-(3-phenylpropyl)-aminopurine (**21**), 6-(4-phenylbutyl)-aminopurine (**22**) (Nishikawa et al., 1986). The initial substrates 2,6-dichloropurine, 8-bromo-6-benzylaminopurine, 2-amino-6-chloropurine were obtained according to ref. (Steklov et al., 2011). 2-Fluoro-6-chloropurine was obtained from 2-amino-6-chloropurine according to ref. (Yu Lin et al., 2010). 8-Bromo-2-fluoro-6-benzylaminopurine and 8-bromo-2-chloro-6-benzylaminopurine were obtained by a bromination of 2-fluoro-6-benzylaminopurine and 2-chloro-6-benzylaminopurine respectively according to ref. (Steklov et al., 2011). *N*^6^-(β-Naphthylmethyl)adenosine was obtained according to ref. (Orlov et al., 2017). In this work we used commercially available 6-benzylthiopurine (**15**), 2-amino-6-benzyloxypurine (**16**) and 6-benzoylaminopurine (**25**) (Sigma). ^1^H NMR and MS data are presented in full in Supplementary information.

**9-methyl-6-benzylaminopurine (2)**. To the mixture of BA (200 mg, 0.89 mmol) and К_2_СО_3_ (250 mg, 1.81 mmol) in DMF (2 ml) MeI (0.07 ml, 1.12 mmol) was added in one portion and the mixture was left to stirred at room temp. The reaction was monitored by TLC (CH_2_Cl_2_:EtOH – 96:4). After 16 h about 80% conversion of initial BA was observed. The additional portions of reagents (К_2_СО_3_, MeI) in above indicated quantities were added to achieve full conversion of BA. After 1 h the reaction mixture was diluted with ethyl acetate (10 ml) and washed with brine (2×20 ml). The organic layer was separated, dried over anhydrous Na_2_SO_4_ and evaporated in vacuum. The residue was purified by column chromatography on silica gel. The product was eluted with CH_2_Cl_2_:EtOH = 97:3 mixture. Yield of *N*^6^-benzyl-9-methyl-adenine was 180 mg (85%) as white powder. ^1^H NMR (400 MHz, CDCl_3_) δ = 8.42 (s, 1H, H2), 7.62 (s, 1H, H8), 7.45–7.10 (m, 5H, Ph), 6.37 (br s, 1H, *N*^6^H), 4.89 (br s, 2H, *N*^6^CH_2_), 3.80 (s, 3H, Me). HRMS: *m/z* [M–H]^−^ calculated 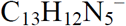 238.1087, found 238.1091; *m/z* [M+Cl]^−^ calculated 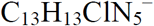 274.0854, found 274.0862.

**9-ethyl-6-benzylaminopurine (3)**. To the mixture of BA (200 mg, 0.89 mmol) and К_2_СО_3_ (500 mg, 3.62 mmol) in DMF (2 ml) (EtO_2_)SO_2_ (0.15 ml, 1.145 mmol) was added in one portion and the mixture was left to stirred at room temp. The reaction was monitored by TLC (CH_2_Cl_2_:EtOH – 96:4). After 16 h about 90% conversion of initial BA was observed. The distilled water (20 ml) was added and the mixture was left to stirred for 1 h. After 1 h the mixture was diluted with ethyl acetate (50 ml) and extracted. The organic layer was separated, washed with brine (2×20 ml), dried over anhydrous Na_2_SO_4_ and evaporated in vacuum. The residue was purified by column chromatography on silica gel. The product was eluted with CH_2_Cl_2_:EtOH = 97:3 mixture. Yield of *N*^6^-benzyl-9-ethyl-adenine was 159 mg (70%) as white powder. ^1^H NMR (400 MHz, CDCl_3_) δ = 8.42 (s, 1H, H2), 7.72 (s, 1H, H8), 7.42–7.23 (m, 5H, Ph), 6.24 (br s, 1H, *N*^6^H), 4.90 (br s, 2H, *N*^6^CH_2_), 4.24 (q, 2H, *J*_CH2-CH3_ = 7.3 Hz, CH_2_CH_3_), 1.53 (t, 3H, *J*_CH2-CH3_ = 7.3 Hz, CH_2_C***H_3_***). HRMS: *m/z* [M–H]^−^ calculated C_14_H_14_N_5_^−^ 252.1244, found 252.1242; *m/z* [M+Cl]^−^ calculated C_14_H_15_ClN_5_^−^ 288.1010, found 288.1011.

**9-isopropyl-6-benzylaminopurine (4)**. Following the procedure for preparation of 9-ethyl-6-benzylaminopurine, the reaction of BA (200 mg, 0.89 mmol) with i-PrBr (0.21 ml, 2.24 mmol) in the presence of К_2_СО_3_ (500 mg, 3.62 mmol) in DMF (2 ml) during 16 h at room temp. gave 9-isopropyl-6-benzylaminopurine as a white powder. Yield 182 mg (77%). ^1^H NMR (400 MHz, CDCl_3_) δ = 8.42 (s, 1H, H2), 7.75 (s, 1H, H8), 7.45–7.20 (m, 5H, Ph), 6.28 (br s, 1H, *N*^6^H), 4.90 (br s, 2H, *N*^6^CH_2_), 4.84 (g, 2H, *J*CH-CH3 = 6.8 Hz, C***H***(CH_3_)_2_), 1.61 (s, 3H, Me), 1.59 (s, 3H, Me). HRMS: *m/z* [M–H]^−^ calculated C_15_H_16_N_5_^−^ 266.1400, found 266.1405; *m/z* [M+Cl]^−^ calculated C_15_H_17_ClN_5_^−^ 302.1167, found 302.1170.

**6-(benzylamino)-9-methyl-7,9-dihydro-purin-8-one (5)**. To the mixture of 9-methyl-6-benzylaminopurine (400 mg, 1.67 mmol) and AcONa×3Н_2_О (1.1 g, 8.08 mmol) in АсОН (4 ml) Br^2^ (0.35 ml, 6.83 mmol) was added in one portion. The mixture was stirred at room temp. for 3.5 h. During the reaction, the initial nucleoside was dissolve. The reaction was monitored by TLC (CH_2_Cl_2_:EtOH – 95:5). The reaction mixture was neutralized with 10% aqueous sodium bicarbonate (80 ml) and washed with methylene chloride (2×50 ml). The organic layer was separated, dried over anhydrous Na_2_SO_4_ and evaporated in vacuum. The residue was purified by column chromatography on silica gel. The product was eluted with CH_2_Cl_2_:EtOAc mixture with gradient from 3:1 to 1:1. Yield of 9-methyl-8-bromo-6-benzylaminopurine was 330 mg (62%) as white powder. R*_f_* = 0.23 (CH_2_Cl_2_ – EtOH, 95:5). 16 h at room temp. gave 6-benzyl-9-isopropyl-adenine as a white powder. Yield 182 mg (77%). ^1^H NMR (400 MHz, CDCl_3_) δ = 8.38 (s, 1H, H2), 7.40–7.20 (m, 5H, Ph), 6.04 (br s, 1H, *N*^6^H), 4.86 (br s, 2H, *N*^6^CH_2_), 3.77 (s, 3H, Me).

The mixture of 9-methyl-8-bromo-6-benzylaminopurine (150 mg, 0.471 mmol) and AcONa×3Н_2_О (640 mg, 4.7 mmol) in АсОН (1.5 ml) was stirred at 120 °C. The reaction was monitored by TLC (CH_2_Cl_2_:EtOH – 95:5). After 16 h the reaction mixture was cooled, neutralized with 10% aqueous sodium bicarbonate (60 ml) and washed with methylene chloride (2×50 ml). The organic layer was separated, dried over anhydrous Na_2_SO_4_ and evaporated in vacuum. Yield of 6-(benzylamino)-9-methyl-7,9-dihydro-purin-8-one (**5**) was 103 mg (85%) as white powder. R*_f_* = 0.23 (CH_2_Cl_2_ – EtOH, 95:5). ^1^H NMR (400 MHz, CDCl_3_) δ = 10.62 (s, 1H, 8-OH), 8.30 (s, 1H, H2), 7.50–7.20 (m, 5H, Ph), 6.68 (br s, 1H, *N*^6^H), 4.84 (br s, 2H, *N*^6^CH_2_), 2.98 (s, 3H, Me). HRMS: *m/z* [M–H]^−^ calculated C_13_H_12_N_5_O^−^ 254.1036, found 254.1037; *m/z* [M+Cl]^−^ calculated C_12_H_13_ClN_5_O^−^ 290.0803, found 290.0803.

**6-(benzylamino)-7,9-dihydro-purin-8-one (6)**. The mixture of 8-bromo-6-benzylaminopurine (169 mg, 0.556 mmol) and KOH (229 mg, 4.082 mmol) in 5 ml of acetic acid was refluxed during 12 h. Then the reaction mixture was evaporated in vacuum and the residue was crystallized in cold water, filtered and washed with cold water. The precipitate was dissolved under heating in ethanol (120 ml), mixed with small amount of silica gel and evaporated in vacuum to dryness. The mixture was applied to chromatographic column on silica gel, using CH_2_Cl_2_:EtOH gradient, increasing in polarity from CH_2_Cl_2_ to CH_2_Cl_2_:EtOH (92:8). Yield of 6-(benzylamino)-7,9-dihydro-purin-8-one (**6**) was 76 mg (57%). R_f_ 0.08 (CH_2_Cl_2_:EtOH – 95:5). ^1^H NMR (400 MHz, DMSO-*d*_6_): δ = 11.27 (br s, 1H, N9H), 9.87 (br s, 1H, 8-OH), 8.04 (s, 1H, H2), 7.40–7.20 (m, 5H, Ph), 6.87 (br s, 1H, *N*^6^H), 4.65 (br s, 2H, *N*^6^CH_2_). HRMS: *m/z* [M–H]^−^ calculated C_12_H_10_N_5_O^−^ 240.0880, found 240.0881; *m/z* [M+Cl]^−^ calculated C_12_H_11_ClN_5_O^−^ 276.0647, found 276.0649.

**6-(benzylamino)-7,9-dihydro-purine-8-thione (7)**. The mixture of 8-bromo-6-benzylaminopurine (221 mg, 0.727 mmol) and thiourea (89 mg, 1.169 mmol) in the mixture of *n*-BuOH and DMF (1:1) (8.8 ml) was stirred at 150 °C during 5 h. Then the mixture was cooled to the room temp., diluted with ethyl acetate (20 ml) and washed successively with water (3×15 ml) and brine (15 ml). The organic layer was separated, dried over anhydrous Na_2_SO_4_, filtered and evaporated in vacuum. The residue was purified by column chromatography on silica gel, using CH_2_Cl_2_:EtOH gradient, increasing in polarity from CH_2_Cl_2_ to CH_2_Cl_2_:EtOH (92:8). Yield of 6-(benzylamino)-7,9-dihydro-purine-8-thione (**7**) was 51 mg (27%). R_f_ 0.31 (CH_2_Cl_2_:EtOH – 95:5). ^1^H NMR (400 MHz, DMSO-*d*_6_): δ = 13.07 (br s, 1H, N9H), 9.87 (br s, 1H, 8-SH), 8.16 (s, 1H, H2), 7.40–7.15 (m, 5H, Ph), 7.14 (br s, 1H, *N*^6^H), 4.69 (br s, 2H, *N*^6^CH_2_). HRMS: *m/z* [M–H]^−^ calculated C_12_H_10_N_5_S^−^ 256.0651, found 256.0655; *m/z* [M+Cl]^−^ calculated C_12_H_11_ClN_5_S^−^ 292.0418, found 292.0423.

**6-(benzylamino)-2-fluoro-7,9-dihydro-purine-8-thione (8)**. The mixture of 8-bromo-2-fluoro-6-benzylaminopurine (206 mg, 0.640 mmol) and thiourea (70 mg, 0.920 mmol) in 10 ml of *n*-BuOH was stirred at 150 °C during 6 h. Then the reaction mixture was evaporated in vacuum and the residue was crystallized in cold water, filtered and washed with cold water. The precipitate was dissolved in dichloromethane and purified by column chromatography on silica gel, using CH_2_Cl_2_:EtOH gradient, increasing in polarity from CH_2_Cl_2_ to CH_2_Cl_2_:EtOH (95:5). Yield of 6-(benzylamino)-2-fluoro-7,9-dihydro-purine-8-thione (**8**) was 63 mg (36%). R_f_ 0.31 (CH_2_Cl_2_:EtOH – 95:5). ^1^H NMR (400 MHz, DMSO-*d*_6_): δ = 13.24 (br s, 1H, N9H), 11.89 (br s, 1H, 8-SH), 7.56 (br s, 1H, *N*^6^H), 7.40–7.25 (m, 5H, Ph), 4.64 (br s, 2H, *N*^6^CH_2_). HRMS: *m/z* [M–H]^−^ calculated C_12_H_9_FN_5_S^−^ 274.0557, found 274.0562.

**6-(benzylamino)-2-chloro-7,9-dihydro-purine-8-thione (9)**. Following the procedure for preparation of 6-(benzylamino)-2-fluoro-7,9-dihydro-purine-8-thione (**8**), the reaction of 8-bromo-2-chloro-6-benzylaminopurine (163 mg, 0.481 mmol) with thiourea (54 mg, 0.709 mmol) in 10 ml of *n*-BuOH during 8 h at 150 °C gave 6-(benzylamino)-2-chloro-7,9-dihydro-purine-8-thione (**9**) as a white powder. Yield 52 mg (37%). R_f_ 0.31 (CH_2_Cl_2_:EtOH – 95:5). ^1^H NMR (400 MHz, DMSO-*d*_6_): δ = 13.25 (br s, 1H, N9H), 11.93 (br s, 1H, 8-SH), 7.46 (br s, 1H, *N*^6^H), 7.40–7.25 (m, 5H, Ph), 4.63 (br s, 2H, *N*^6^CH_2_). HRMS: *m/z* [M–H]^−^ calculated C_12_H_9_ClN_5_S^−^ 290.0262, found 290.0265; *m/z* [M+Cl]^−^ calculated C_12_H_10_Cl_2_N_5_S^−^ 326.0028, found 326.0018.

**2-fluoro-6-benzylaminopurine (11)**. To the solution of 2-fluoro-6-chloropurine (949 mg, 5.5 mmol) in 20 ml of *n*-BuOH DIPEA (1.13 ml, 7.76 mmol) was added in one portion and the mixture was left to stirred at room temperature for 5 min. Then benzylamine (0.613 ml, 5.61 mmol) was added in one portion and the mixture was left to stirred at 65 °C for 5 h. Then the reaction mixture was evaporated in vacuum and the residue was crystallized in cold water, filtered and washed with cold water. The precipitate was dissolved in dichloromethane and purified by column chromatography on silica gel, using CH_2_Cl_2_:EtOH gradient, increasing in polarity from CH_2_Cl_2_ to CH_2_Cl_2_:EtOH (92:8). Yield of 2-fluoro-6-benzylaminopurine (**11**) was 1204 mg (90%). R_f_ 0.21 (ethyl acetate:hexane – 2:1). ^1^H NMR (400 MHz, DMSO-*d*_6_): δ = 13.02 (br s, 1H, 9-NH), 8.74 (br s, 1H, *N*^6^H), 8.08 (s, 1H, H8), 7.40–7.20 (m, 5H, Ph), 4.64 (br s, 2H, *N*^6^CH_2_). HRMS: *m/z* [M–H]^−^ calculated C_12_H_9_FN_5_^−^ 242.0836, found 242.0836; *m/z* [M+Cl]^−^ calculated C_12_H_10_FClN_5_^−^ 278.0603, found 278.0606.

**2-chloro-6-(3-fluorobenzyl)-aminopurine (12)**. Following the procedure for preparation of 2-fluoro-6-benzylaminopurine (**11**), the reaction of 2,6-dichloropurine (143 mg, 0.757 mmol) with 3-fluorobenzylamine (0.095 ml, 0.795 mmol) in the presence of DIPEA (0.195 ml, 1.176 mmol) in 3.5 ml of *n*-BuOH during 4 h at 85 °C gave 2-chloro-6-(3-fluorobenzyl)-aminopurine (**12**) as a white powder. Yield 207 mg (98%). R_f_ 0.52 (CH_2_Cl_2_:EtOH – 92:8). ^1^H NMR (400 MHz, DMSO-*d*_6_): δ = 13.10 (br s, 1H, 9-NH), 8.73 (br s, 1H, *N*^6^H), 8.15 (s, 1H, H8), 7.40–7.00 (m, 4H, Ph), 4.66 (br s, 2H, *N*^6^CH_2_). HRMS: *m/z* [M–H]^−^ calculated C_12_H_8_ClFN_5_^−^ 276.0447, found 276.0447; *m/z* [M+Cl]^−^ calculated C_12_H_9_Cl_2_FN_5_^−^ 312.0214, found 312.0209.

**2-amino-6-(2-phenylethoxy)-purine (17)**. The mixture of 2-amino-6-chloropurine (1000 mg, 5.9 mmol) and DABCO (3310 mg, 29.5 mmol) in DMSO (6 ml) was left at room temp. overnight. Then isopropanol (10 ml) was added, the resulting precipitate was filtered, washed with isopropanol (3×10 ml) and dried in vacuum over P_2_O_5_. The yield of 1-(2-aminopurin-6-yl)-1,4-diazabicyclo[2.2.2]octan-1-ium chloride was 1441 mg (87%) as white powder. The mixture of 1-(2-aminopurin-6-yl)-1,4-diazabicyclo[2.2.2]octan-1-ium chloride (300 mg, 1.06 mmol) and t-BuOK (357 mg, 3.18 mmol) in 2-phenylethanol (6.1 ml, 50.56 mmol) was stirred at room temp. during 72 h. The reaction was monitored by TLC (CH_2_Cl_2_:EtOH – 90:10). After 72 h the reaction mixture was neutralized with acetic acid (0.12 ml, 2.127 mmol). The mixture without evaporation in vacuum was applied to chromatographic column on silica gel. The product was eluted using CH_2_Cl_2_:EtOH gradient, increasing in polarity from CH_2_Cl_2_ to CH_2_Cl_2_:EtOH (90:10). Yield of 2-amino-6-(2-phenylethoxy)-purine (**17**) was 117 mg (43%) as a white powder. R_f_ 0.37 (CH_2_Cl_2_:EtOH – 90:10). ^1^H NMR (400 MHz, DMSO-*d*_6_): δ = 12.36 (br s, 1H, 9-NH, 7.78 (s, 1H, H8), 7.37–7.20 (m, 5H, Ph), 6.19 (br s, 2H, 2-NH_2_), 4.60 (t, 2H, *J*_CH2CH2_ = 7.2 Hz, OC***H*_2_**CH_2_Ph), 3.08 (t, 2H, *J*_CH2CH2_ = 7.2 Hz, OCH_2_C***H*_2_**Ph). HRMS: *m/z* [M–H]^−^ calculated C_13_H_12_N_5_O^−^ 254.1036, found 254.1041; *m/z* [M+Cl]^−^ calculated C_13_H_13_ClN_5_O^−^ 290.0803, found 290.0809.

**2-amino-6-(3-phenylpropoxy)-purine (18)**. Following the procedure for preparation of 2-amino-6-(2-phenylethoxy)-purine (**17**), the reaction of 1-(2-aminopurin-6-yl)-1,4-diazabicyclo[2.2.2]octan-1-ium chloride (200 mg, 0.709 mmol) with 3-phenylpropan-1-ol (4.6 ml, 33.8 mmol) in the presence of t-BuOK (239 mg, 2.127 mmol) during 72 h at room temp. gave 2-amino-6-(3-phenylpropoxy)-purine (**18**) as a white powder. Yield 127 mg (67%). R_f_ 0.33 (CH_2_Cl_2_:EtOH – 92:8). ^1^H NMR (400 MHz, DMSO-*d*_6_): δ = 12.36 (br s, 1H, 9-NH, 7.79 (s, 1H, H8), 7.32–7.15 (m, 5H, Ph), 6.16 (br s, 2H, 2-NH_2_), 4.39 (t, 2H, *J*_CH2CH2_ = 6.6 Hz, OCH_2_CH_2_C***H*_2_**Ph), 3.08 (t, 2H, *J*_CH2CH2_ = 7.4 Hz, OC***H*_2_**CH_2_CH_2_Ph), 2.08 (tt, 2H, *J*_CH2CH2_ = 7.4 Hz, *J*_CH2CH2_ = 6.6 Hz, OCH_2_C***H*_2_**CH_2_Ph). HRMS: *m/z* [M–H]^−^ calculated C_14_H_14_N_5_O^−^ 268.1193, found 268.1200; *m/z* [M+Cl]^−^ calculated C_14_H_15_ClN_5_O^−^ 304.0960, found 304.0962.

**2-chloro-6-furfurylaminopurine (23)**. To the suspension of 2,6-dichloropurine (205 mg, 1.085 mmol) in 3.2 ml of 2-propanol furfurylamine (0.341 ml, 3.827 mmol) was added in one portion and the mixture was left to stirred at 90 °С for 3.5 h. Then the reaction mixture was evaporated in vacuum and the residue was crystallized in cold water (9 ml), filtered and washed with cold water and 2-propanol. The product was recrystallized in ethanol (3 ml). Yield of 2-chloro-6-furfurylaminopurine (**23**) was 215 mg (79%). R_f_ 0.16 (ethyl acetate:hexane – 2:1). ^1^H NMR (400 MHz, DMSO-*d*_6_): δ = 13.08 (br s, 1H, 9-NH), 8.54 (br s, 1H, *N*^6^H), 8.14 (s, 1H, H8), 7.56 (s, 1H, furan), 6.38 (s, 1H, furan), 6.28 (s, 1H, furan), 4.64 (br s, 2H, *N*^6^CH_2_). HRMS: *m/z* [M–H]^−^ calculated C_10_H_7_ClN_5_O^−^ 248.0334, found 248.0331; *m/z* [M+Cl]^−^ calculated C_10_H_8_Cl_2_N_5_O^−^ 284.0100, found 284.0100.

**2-chloro-6-(1-phenylethyl)-aminopurine (24)**. Following the procedure for preparation of 2-chloro-6-furfurylaminopurine (**23**), the reaction of 2,6-dichloropurine (194 mg, 1.026 mmol) with α-phenylethylamine (0.156 ml, 1.232 mmol) in the presence of DIPEA (0.211 mg, 1.233 mmol) in 3 ml of 2-propanol during 6 h at 90 °С gave 2-chloro-6-(1-phenylethyl)-aminopurine (**24**) as a white powder. The product was recrystallized from 2-propanol. Yield 196 mg (70%). R_f_ 0.16 (ethyl acetate:hexane – 2:1). ^1^H NMR (400 MHz, DMSO-*d*_6_): δ = 13.04 (br s, 1H, 9-NH), 8.59 (br s, 1H, *N*^6^H), 8.11 (s, 1H, H8), 7.48–7.15 (m, 5H, Ph), 5.40 (br s, 1H, C***H***MePh), 1.54 (d, 3H, *J*CHCH_3_ = 6.7 Hz, Me). HRMS: *m/z* [M–H]^−^ calculated C_13_H_11_ClN_5_^−^ 272.0697, found 272.0699; *m/z* [M+Cl]^−^ calculated C_13_H_12_Cl_2_N_5_^−^ 308.0464, found 308.0457.

**6-(β-naphthylmethyl)-aminopurine (26)**. The solution of *N*^6^-(β-naphthylmethyl)-adenosine (101 mg, 0.248 mmol) in 0.5M НСl (3.5 ml, 1.77 mmol) was stirred at 100 °С. The reaction was monitored by TLC (acetonitrile - 25% ammonia, 9:1). After 2 h when the traces of initial nucleoside had disappeared, the reaction was cooled and neutralized to pH 7 with concentrated NH_3_. The resulting precipitate was filtered, washed with cold water (3×5 ml) and dried in vacuum over P_2_O_5_. The yield of 6-(β-naphthylmethyl)-aminopurine (**26**) was 65 mg (94%) as white powder. R_f_ = 0.40 (acetonitrile - 25% ammonia, 9:1). mp = 245–247 °C. ^1^H NMR (400 MHz, DMSO-*d*_6_): δ = 4.96 (br s, 2H, *N*^6^CH_2_), 7.43–7.58 (m, 3H, naphthalene), 7.82–7.92 (m, 5H, *N*^6^H, naphthalene), 8.31 (s, 1H, H2), 8.38 (s, 1H, H8), 9.00 (br s, 1H, 9-NH). HRMS: *m/z* [M−H]^−^ calculated C_16_H_12_N_5_− 274.1087, found 274.1089; *m/z* [M+Cl]^−^ calculated C_16_H_13_N_5_Cl− 310.0854, found 310.0848.

### 6.5. Molecular modeling

Previously published homology model of AHK3 based on 3T4L template was used (Lomin et al., 2015), and the same modeling protocol was used for AHK2. Alignment used for modeling is shown in Supplementary Fig. 1.

#### 6.5.1. Binding mode analysis

Structure preparation, docking study and the postprocessing of the obtained results were performed using OpenEye Toolkits (OpenEye Toolkits 2017.Feb.1 OpenEye Scientific Software, Santa Fe, NM. http://www.eyesopen.com). First, the reasonable tautomers were generated using the QuacPac Toolkit (Quacpac Toolkit 2017.Feb.1 OpenEye Scientific Software, Santa Fe, NM. http://www.eyesopen.com), which in most cases appeared to be the most favorable with the following parameter set: *CarbonHybridization* = *False*, *MaxTautomericAtoms* = 30, *MaxZoneSize* = 50, *RankTautomers* = *True*, *MaxTautomersGenerated* = 256. Then the exhaustive enantiomer and conformation generation was carried out with the Omega Toolkit (Hawkins et al. 2010) (maximum number of conformers of 10000 with default rmsd threshold of 0.5 Å) using MMFF94s force field (Halgren 1999). Energy window value was set to 10 kcal/mol, hydrogen locations for -OH, -NH_2_, -SH were sampled. Receptor structure preparation was performed with *make_receptor* program, which is a part of OEDocking program suite. Both HYBRID (McGann 2012) and FRED (McGann 2011) calculations were carried out with OEDocking toolkit (OEDocking Toolkit 2017.Feb.1 OpenEye Scientific Software, Santa Fe, NM. http://www.eyesopen.com) with high resolution. The obtained poses were scored with Chemgauss4 scoring function and optimized using SZYBKI toolkit (Szybki Toolkit 2017.Feb.1 OpenEye Scientific Software, Santa Fe, NM. http://www.eyesopen.com) in Cartesian coordinates with AM1BCC (Jakalian et al. 2002) charges on ligand atoms and MMFF94s force field using BGFS optimiser (10000 steps and tolerance = 10^−6^). Protein residues inside the sphere of 10 Å of the ligand atoms were treated as flexible.

#### 6.5.2. Quantum Chemical Calculations

All quantum chemical calculations were performed with ORCA 3.0.2 (Neese 2012) using Ahlrichs polarized triple-zeta basis set (def2-TZVPP) (Weigend and Ahlrichs 2005) with tight convergence criteria for both SCF and geometry optimization. Starting geometries were manually created from the *N*^6^-benzyladenine conformation observed in 3T4K structure using Avogadro (Hanwell et al. 2012). Single point energies were calculated with Hartree-Fock, MP2 and hybrid DFT method PBE0 (Perdew and Ernzerhof 1996), which performs well on both energy end electron density calculations.

#### 6.5.3. Artwork

The 3D depictions of the binding site were created with VIDA 4.3.0 (OpenEye Scientific Software, Santa Fe, NM. http://www.eyesopen.com). 2D binding site depictions were generated using Grapheme Toolkit 2017.Feb.1 (OpenEye Scientific Software, Santa Fe, NM. http://www.eyesopen.com). Plots were created in Rstudio (http://www.rstudio.com/) using ggplot2 package (https://cran.r-project.org/web/packages/ggplot2/index.html)

### 6.6. Statistics

All experimental data are means of two independent determinations with s.e. The experiments were carried out in 2–3 biological replications. Correlation coefficients were determined according to Spearman's algorithm using SigmaPlot program.

## Acknowledgments

We thank Dr. K.M. Polyakov (IMB RAS) for valuable suggestions. This study was supported by a grant No 17–04–00969 from the Russian Foundation for Basic Research, and partially by a program of the Presidium of the Russian Academy of Sciences "Molecular and Cell Biology". The preparation of cytokinin derivatives 1–26 was supported by the Russian Science Foundation, grant No 16–14–00178. Free academic licenses for OpenEye software were kindly provided by OpenEye Scientific Software Inc. to the Laboratory of Vladimir A. Palyulin, represented here by D.S.K. and D.I.O.

**Appendix A. Supplementary data**.

## References

Biddington N.L., Thomas T.H., 1973. A modified *Amaranthus* betacyanin bioassay for the rapid determination of cytokinins in plant extracts. Planta 111, 183–186.

Brenner W.G., Romanov G.A., Köllmer I., Bürkle L., Schmülling T., 2005. Immediate-early and delayed cytokinin response genes of *Arabidopsis thaliana* identified by genome-wide expression profiling reveal novel cytokinin-sensitive processes and suggest cytokinin action through transcriptional cascades. Plant J. 44, 314–333.

Clark M., Cramer R.D., Van Opdenbosch N., 1989. Validation of the general purpose Tripos 5.2 force field. J. Comput. Chem. 10, 982–1012.

Doležal K., Popa I., Kryštof V., Spíchal L., Fojtíková M., Holub J., Lenobel R., Schmülling T., Strnad M., 2006. Preparation and biological activity of 6-benzylaminopurine derivatives in plants and human cancer cells. Bioorg Med Chem. 14, 875–884.

Doležal K., Popa I., Hauserová E., Spíchal L., Chakrabarty K., Novák O., Kryštof V., Voller J., Holub J., Strnad M., 2007. Preparation, biological activity and endogenous occurrence of N6-benzyladenosines. Bioorg Med Chem. 15, 3737–3747.

Halgren T.A., 1999. MMFF VI. MMFF94s option for energy minimization studies. J Comput Chem. 20, 720–729.

Hanwell M.D., Curtis D.E., Lonie D.C., Vandermeersch T., Zurek E., Hutchison G.R., 2012. Avogadro: an advanced semantic chemical editor, visualization, and analysis platform. J Cheminf. 4, 17.

Hawkins P.C.D., Skillman A.G., Warren G.L., Ellingson B.A., Stahl M.T., 2010. Conformer Generation with OMEGA: Algorithm and Validation Using High Quality Structures from the Protein Databank and the Cambridge Structural Database. J Chem Inf Model. 50, 572–584.

Heyl A., Riefler M., Romanov G.A., Schmülling T., 2012. Properties, functions, and evolution of cytokinin receptors. Eur J Cell Biol. 91, 246–256.

Hothorn M., Dabi T., Chory J., 2011. Structural basis for cytokinin recognition by *Arabidopsis thaliana* histidine kinase 4. Nature Chem. Biol. 7, 766–768.

Inoue T., Higuchi M., Hashimoto Y., Seki M., Kobayashi M., Kato T., Tabata S., Shinozaki K., Kakimoto T., 2001. Identification of CRE1 as a cytokinin receptor from Arabidopsis. Nature 409, 1060–1063.

Jakalian A., Jack D.B., Bayly C.I., 2002. Fast, efficient generation of high-quality atomic charges. AM1-BCC model: II. Parameterization and validation. J Comput Chem. 23, 1623–1641.

Kakimoto T., 2003. Perception and signal transduction of cytokinins. Annu. Rev. Plant Biol. 54, 605–627.

Kieber J.J., Schaller G.E., 2014. Cytokinins. The Arabidopsis Book 12, e0168.

Kolyachkina S.V., Tararov V.I., Alexeev C.S., Krivosheev D.M., Romanov G.A., Stepanova E.V., Slomko E.S., Andrey N. Inshakov A.N., Mikhailov S.N., 2011. N6-Substituted adenosines. Cytokinin and antitumor activities. Collect. Czech. Chem. Commun. 76, 1361–1378.

Krivosheev D.M., Kolyachkina S.V., Mikhailov S.N., Tararov V.I., Vanyushin B.F., Romanov G.A., 2012. N^6^-(benzyloxymethyl)adenosine is a novel anticytokinin, an antagonist of cytokinin receptor CRE1/AHK4 of Arabidopsis. Dokl Biochem Biophys. 444, 178–181.

Laloue M., Fox J.E., 1989. Cytokinin oxidase from wheat: partial purification and general properties. Plant Physiol. 90, 899–906.

Larkin M.A., Blackshields G., Brown N.P., Chenna R., McGettigan P.A., McWilliam H., Valentin F., Wallace I.M., Wilm A., Lopez R., Thompson J.D., Gibson T.J., Higgins D.G., 2007. Clustal W and Clustal X version 2.0. Bioinformatics 23, 2947–2948.

Laskowski R.A., MacArthur M.W., Moss D.S., Thornton J.M., 1993. PROCHECK - a program to check the stereochemical quality of protein structures. J. App. Cryst. 26, 283–291.

Lomin S.N., Krivosheev D.M., Steklov M.Y., Osolodkin D.I., Romanov G.A., 2012. Receptor properties and features of cytokinin signaling. Acta Naturae 4(3), 31–45.

Lomin S.N., Krivosheev D.M., Steklov M.Yu., Arkhipov D.V., Osolodkin D.I., Schmülling T., Romanov G.A., 2015. Plant membrane assays with cytokinin receptors underpin the unique role of free cytokinin bases as biologically active ligands. J. Exp. Bot. 66, 1851–1863.

Lomin S.N., Yonekura-Sakakibara K., Romanov G.A., Sakakibara H., 2011. Ligand-binding properties and subcellular localization of maize cytokinin receptors. J. Exp. Bot. 62, 5149–5159.

Matsubara S., 1990. Structure-activity relationships of cytokinins. Crit. Rev. Plant Sci. 9, 17–57.

McGann M., 2011. FRED Pose Prediction and Virtual Screening Accuracy. J Chem Inf Model. 51, 578–596.

McGann M., 2012. FRED and HYBRID docking performance on standardized datasets. J Comput Aided Mol Des. 26, 897–906.

Miller C.O., Skoog F., von Saltza N.M., Strong F.M., 1955. Kinetin, a cell division Factor from deoxyribonucleic acid. J. Am. Chem. Soc. 77, 1329–1334.

Mizuno T., Yamashino T., 2010. Biochemical characterization of plant hormone cytokinin-receptor histidine kinases using microorganisms. In: Methods in Enzymology; Vol. 471, Chapter 18. Elsevier Inc. 335–356.

Neese F., 2012. The ORCA program system. Wiley Interdisciplinary Rev. Comp Mol Sci. 2, 73–78.

Nisler J., Zatloukal M., Popa L, Doležal K., Strnad M., Spíchal L., 2010. Cytokinin receptor antagonists derived from 6-benzylaminopurine. Phytochemistry 71, 823–830.

Nishikawa S., Kumazawa Z., Kashimura N., Nishkimi Y., Uemura S., 1986. Alternating dependency of cytokinin activity on the number of methylene units in ω-phenylalkyl derivatives of some purine cytokinins and 4-substituted pyrido[3,4-d]pyrimidine. Agric. Biol. Chem. 50, 2243–2249.

Orlov A.A., Drenichev M.S., Oslovsky V.E., Kurochkin N.N., Solyev P.N., Kozlovskaya L.I., Palyulin V.A., Karganova G.G., Mikhailov S.N., Osolodkin D.I., 2017. New tools in nucleoside toolbox of tick-borne encephalitis virus reproduction inhibitors. Bioorg. Med. Chem. Lett. 27, 1267–1273.

Perdew J.P., Ernzerhof M., 1996. Rationale for mixing exact exchange with density functional approximations. J Chem Phys. 105, 9982–9985.

Plíhal O., Szüčová L., Galuszka P., 2013. N9-substituted aromatic cytokinins with negligible side effects on root development are an emerging tool for in vitro culturing. Plant Signal. Behav. 8, e24392.

Podlešáková K., Zalabák D., Cudejková M., Plíhal O., Szüčová L., Doležal K., Spíchal L., Strnad M., Galuszka P., 2012. Novel cytokinin derivatives do not show negative effects on root growth and proliferation in submicromolar range. PLoS One 7, e39293.

Rashotte A.M., Carson S.D.B., To., J.P.C., Kieber, J.J., 2003. Expression profiling of cytokinin action in Arabidopsis. Plant Physiol. 132, 1998–2011.

Riefler M., Novak O., Strnad M., Schmülling T, 2006. *Arabidopsis* cytokinin receptor mutants reveal functions in shoot growth, leaf senescence, seed size, germination, root development, and cytokinin metabolism. The Plant Cell 18, 40–54.

Romanov G.A., 2009. How do cytokinins affect the cell? Russ. J. Plant Physiol. 56, 268–290.

Romanov G.A., Getman I.A., Schmülling T., 2000. Investigation of early cytokinin effects in a rapid Amaranthus seedling test. Plant Growth Regul. 32, 337–344.

Romanov G.A., Lomin S.N., Schmülling T., 2006. Biochemical characteristics and ligand-binding properties of *Arabidopsis* cytokinin receptor AHK3 compared to CRE1/AHK4 as revealed by a direct binding assay. J. Exp. Bot. 57, 4051–4058.

Romanov G.A., Lomin S.N., 2009. Hormone-binding assay using living bacteria expressing eukaryotic receptors. Plant Hormones: Methods and Protocols, 2nd Edition, Methods in Molecular Biology, Humana Press 495, 111–120.

Romanov G.A., Spíchal L., Lomin S.N., Strnad M., Schmülling T., 2005. A live cell hormone-binding assay on transgenic bacteria expressing a eukaryotic receptor protein. Anal. Biochem. 347, 129–134.

Sakakibara, H., 2006. Cytokinins: activity, biosynthesis, and translocation. Annu. Rev. Plant Biol. 57, 431–449.

Sidorov, G.V., Myasoedov, N.F., Lomin S.N., Romanov G.A., 2015. Synthesis of tritium- and deuterium-labeled isopentenyladenine. Radiochem. 57, 108–110.

Spíchal L., Rakova, N.Y., Riefler M., Mizuno T., Romanov G.A., Strnad M., Schmülling T., 2004. Two cytokinin receptors of *Arabidopsis thaliana*, CRE1/AHK4 and AHK3, differ in their ligand specificity in a bacterial assay. Plant Cell Physiol. 45, 1299–1305.

Spíchal L., Werner T., Popa I., Riefler M., Schmülling T., Strnad M., 2009. The purine derivative PI-55 blocks cytokinin action via receptor inhibition. FEBS J. 276, 244–253.

Steklov M.Yu., Lomin S.N., Osolodkin D.I., Romanov G.A., 2013. Structural basis for cytokinin receptor signaling: an evolutionary approach. Plant Cell Reports 32, 781–793.

Steklov M.Y., Tararov V.I., Romanov G.A., Mikhailov S.N., 2011. Facile synthesis of 8-azido-6-benzylaminopurine. Nucleosides, Nucleotides, Nucleic Acids 30, 503–511.

Stolz A., Riefler M., Lomin S.N., Achazi K., Romanov G.A., Schmülling T., 2011. The specificity of cytokinin signalling in Arabidopsis thaliana is mediated by differing ligand affinities and expression profiles of the receptors. The Plant J. 67, 157–168.

Strnad M., 1997. The aromatic cytokinins. Physiol. Plant. 101, 674–688.

Suzuki T., Miwa K., Ishikawa K., Yamada H., Aiba H., Mizuno T., 2001. The Arabidopsis sensor His-kinase, AHK4, Can Respond to Cytokinins. Plant Cell Physiol. 42, 107–113.

Szücová L., Spíchal L., Dolezal K., Zatloukal M., Greplová J., Galuszka P., Krystof V., Voller J., Popa I., Massino F.J., Jørgensen J.E., Strnad M., 2009. Synthesis, characterization and biological activity of ring-substituted 6-benzylamino-9-tetrahydropyran-2-yl and 9-tetrahydrofuran-2-ylpurine derivatives. Bioorg. Med. Chem. 17, 1938–1947.

Tararov V.I., Kolyachkina S.V., Alexeev C.S., Mikhailov S.N., 2011. N6-Acetyl-2′,3′,5′-tri-O-acetyladenosine; a convenient, ‘Missed Out’ substrate for regioselective N6-alkylations. Synthesis 15, 2483–2489.

Theiler J.B., Leonard N.J., Schmitz R.Y., Skoog F., 1976. Photoaffinity-labeled cytokinins synthesis and biological activity. Plant Physiol. 58, 803–805.

Vylíčilová H., Husičková A., Spíchal L., Srovnal J., Doležal K., Plíhal O., Plíhalová L., 2016. C2-substituted aromatic cytokinin sugar conjugates delay the onset of senescence by maintaining the activity of the photosynthetic apparatus. Phytochemistry 122, 22–33.

Wang J., Wang Q., Zhang L., Fang, H., 2013. Design, synthesis and preliminary biological evaluation of purine-2,6-diamine derivatives as cyclin-dependent kinase (CDK) inhibitors. Chinese J. Chem. 31, 1181–1191.

Weigend F., Ahlrichs R., 2005. Balanced basis sets of split valence, triple zeta valence and quadruple zeta valence quality for H to Rn: Design and assessment of accuracy. Phys Chem Chem Phys. 7, 3297–3305.

Yu Lin H., Xiang L., Ming L., 2010. Synthesis and biological activity of novel 6-substituted purine derivatives. J. Mex. Chem. Soc. 54, 74–78.

Zahajská L., Nisler J., Voller J., Gucký T., Pospíšil T., Spíchal L., Strnad M., 2017. Preparation, characterization and biological activity of C8-substituted cytokinins. Phytochemistry 135, 115–127.

Zatloukal M., Gemrotová M., Doležal K., Havlíček L., Spíchal L., Strnad M., 2008. Novel potent inhibitors of A. thaliana cytokinin oxidase/dehydrogenase. Bioorg Med Chem. 16, 9268–9275.

Zvereva S.D., Romanov G.A., 2000. Reporter genes for plant genetic engineering: characteristics and detection. Russ. J. Plant Physiol. 47, 424–432.

